# The Histone Methyltransferase SETD2 Modulates Oxidative Stress to Attenuate Colonic Inflammation and Tumorigenesis in Mice

**DOI:** 10.1101/2020.07.13.201624

**Authors:** Min Liu, Hanyu Rao, Jing Liu, Xiaoxue Li, Wenxin Feng, Jin Xu, Wei-Qiang Gao, Li Li

**Affiliations:** State Key Laboratory of Oncogenes and Related Genes, Renji-Med X Clinical Stem Cell Research Center, Ren Ji Hospital, School of Medicine and School of Biomedical Engineering, Shanghai Jiao Tong University, Shanghai, 200127, China; School of Biomedical Engineering and Med-X Research Institute, Shanghai Jiao Tong University, Shanghai, China

**Keywords:** IBD, SETD2, Oxidative Stress, ROS, Epithelial Barrier

## Abstract

**BACKGROUND & AIMS:** Inflammatory bowel disease (IBD) is a complex and relapsing inflammatory disease, and patients with IBD exhibit a higher risk of developing colorectal cancer (CRC). Epithelial barrier disruption is one of the major causes of IBD in which epigenetic modulation is pivotal. However, the epigenetic mechanisms underlying the epithelial barrier integrity regulation remain largely unexplored. Here, we investigated how SETD2, an epigenetic modifier, maintains intestinal epithelial homeostasis and attenuates colonic inflammation and tumorigenesis.

**METHODS:** GEO public database and IBD tissues were used to investigate the clinical relevance of SETD2 in IBD. To define a role of SETD2 in the colitis, we generated mice with epithelium-specific deletion of *Setd2* (*Setd2*^*Vil-KO*^ mice). Acute colitis was induced by 2% dextran sodium sulfate (DSS), and colitis-associated CRC was induced by injecting azoxymethane (AOM), followed by three cycles of 2% DSS treatments. Colon tissues were collected from mice and analyzed by histology, immunohistochemistry and immunoblots. Organoids were generated from Setd2^Vil-KO^ and control mice, and were stained with 7-AAD to detect apoptosis. A fluorescent probe, 2′,7′-dichlorodihydrofluorescein diacetate (H2DCFDA), was used to detect the levels of ROS in intestinal epithelial cells (IECs) isolated from the two types of mice. RNA-seq and H3K36me3 ChIP-seq analyses were performed to identify the mis-regulated genes modulated by SETD2. Results were validated in functional rescue experiments by N-acetyl-l-cysteine (NAC) treatment and transgenes expression in IECs.

**RESULTS:** SETD2 expression became decreased in IBD patients and DSS-treated colitis mice. *Setd2*^*Vil-KO*^ mice displayed abnormal loss of mucus-producing goblet cells and antimicrobial peptide (AMP)-producing Paneth cells, and exhibited pre-mature intestinal inflammation development. Consistent with the reduced SETD2 expression in IBD patients, *Setd2*^*Vil-KO*^ mice showed increased susceptibility to DSS-induced colitis, accompanied by more severe epithelial barrier disruption and markedly increased intestinal permeability that subsequently facilitated inflammation-associated CRC. Mechanistically, deletion of *Setd2* resulted in excess reactive oxygen species (ROS), which led to cellular apoptosis and defects in barrier integrity. NAC treatment in *Setd2*^*Vil-KO*^ mice rescued epithelial barrier injury and apoptosis. Importantly, *Setd2* depletion led to excess ROS by directly down-regulating antioxidant genes that inhibit ROS reaction. Moreover, overexpression of antioxidant PRDX6 in *Setd2*^*Vil-KO*^ IECs largely alleviated the overproductions of ROS and improved the cellular survival.

**CONCLUSIONS:** Deficiency of Setd2 specifically in the intestine aggravates epithelial barrier disruption and inflammatory response in colitis via a mechanism dependent on oxidative stress. Thus, our results highlight an epigenetic mechanism by which Setd2 modulates oxidative stress to regulate intestinal epithelial homeostasis. SETD2 might therefore be a pivotal regulator that maintains the homeostasis of the intestinal mucosal barrier.

## Introduction

Inflammatory bowel disease (IBD) is the most commonly diagnosed inflammatory disorder that can be sub-classified into Crohn’s disease (CD) and ulcerative colitis (UC)^1, 2^. Chronic inflammation inevitably results in tissue injury and tumorigenesis, and IBD patients have higher risk of colorectal cancer development^3, 4^. IBD results from a complex interactions between the mucosal barrier, commensal bacteria, and the immune system^5^. The intestinal mucosal barrier consists of intestinal epithelial cells (IECs), which are adhered to each other via epithelial tight junctions, and plays a critical role in protecting the gastrointestinal tract^6^. Defects in the barrier function allow the translocation of luminal pathogens and bacteria into the intestinal lamina propria and trigger chronic inflammation and tissue injury, which occasionally causes dysplasia^7^. IBD have been associated with abnormal intestinal epithelial barrier function^8, 9^. However, the underlying molecular mechanisms of IBD remain largely ambiguous.

Intestinal inflammation is usually accompanied by excessive production of reactive oxygen. High levels of reactive oxygen species (ROS) create oxidative stress within the cells and lead to a loss of homeostasis^10^. Redox imbalance has been proposed as one potential cause factor for IBD^11^. Oxidative stress plays an important role in the development and progression of IBD, and is associated with CRC pathogenesis^12-14^. In the gastrointestinal tract, oxidative stress leads to damages of the intestinal mucosal layer and epithelial cell apoptosis, which results in bacterial invasion in the gut and in turn stimulates the immune response^15-17^. Clinical studies show that the increase of ROS and biomarkers of oxidative injury contributes to tissue damage in IBD^17, 18^. Our previous work has also shown that excess ROS leads to defects in barrier integrity and the subsequent inflammation^19^. In order to resist the oxidative damage, the intestinal epithelium contains an extensive system of antioxidants^20, 21^. Colitis is usually associated with a decrease in the levels of antioxidants in colonic tissues^18^, whereas overexpression of antioxidant enzymes results in attenuation of colitis in mice^16^. Thus, a balance between oxidant and antioxidant mechanisms is necessary to maintain intestinal epithelial homeostasis.

Epigenetic regulation disorder is important in the onset and pathogenesis of IBD^22-24^, which may influence the maintenance of homeostasis in the intestinal epithelium. Studies have shown that DNA methylation is correlated with disease susceptibility and progression in IBD^25, 26^. It is also known that inactive histone modification H3K27 trimethylation regulated by EZH2 promotes the inflammatory response and apoptosis in colitis^27^. Moreover, our previous study has also shown that BRG1, a key epigenetic regulator, is required for the homeostatic maintenance of intestines to prevent inflammation and tumorigenesis^19^. In addition, SETD2, a trimethyltransferase of histone H3 lysine 36 (H3K36), is frequently mutated (17%) in UC samples with a high risk of developing colorectal carcinoma^28^, suggesting that SETD2 plays a potential role in colitis and colitis-associated CRC. SETD2 has also been identified as a common mutation across cancer types, including glioma^29^, renal cell carcinoma^30^, leukemia^31, 32^ and lung cancer^33, 34^. In addition, we have recently also reported that SETD2 is pivotal for bone marrow mesenchymal stem cells differentiation, genomic imprinting and embryonic development, initiation and metastasis of pancreatic cancer, and normal lymphocyte development^35-39^. However, functions of SETD2 in IBD and inflammation-associated CRC remain largely undefined.

To investigate a possible role of SETD2 in IBD, in the present work we generated intestinal epithelium-specific *Setd2*-knockout mice, and found that deletion of *Setd2* in intestinal tissues promotes DSS-induced epithelial damage and subsequent tumorigenesis. Mechanistically, our results highlight SETD2 as a critical epigenetic determinant in the prevention of colitis and tumorigenesis through modulation of oxidative stress.

## Materials and Methods

The full Materials and Methods section is included in the supplementary information.

### Animal Experiments

All animals were bred and maintained at an animal facility under specific pathogen-free conditions. Sex-matched littermates were used in all studies. All studies with mice were carried out by following the Guide for the Care and Use of Laboratory Animals, which was approved by Bioethics Committee at the School of Biomedical Engineering, Shanghai Jiao Tong University. *Setd2*^*Vil-KO*^ mice were generated by crossing *Setd2-floxed* mice with *Villin-Cre* mice. Acute colitis was induced by adding 2% dextran sodium sulphate (DSS; MP Biomedicals) for 5 days, which was then followed by 5 days of regular drinking water. Body weights were monitored daily, and mice were sacrificed 5 days and 10 days after DSS treatment, except when indicated otherwise. Colitis-associated CRC was induced by injecting AOM (10 mg/kg, Sigma), followed by three cycles of 2% DSS treatments.

### Isolation of Intestinal Epithelial Cells and Culture

For RNA and protein extraction analyses, IECs were isolated as follows: mice were sacrificed and colonic specimens were dissected, opened longitudinally and cleared from feces by washing extensively in cold PBS. Colons were cut in small pieces of 1 mm and incubated in 30 mM EDTA solution in PBS at 37°C for 10 min. 10 min later, EDTA solution was replaced with ice-cold PBS and shacked vigorously for 30 s. This process was repeated once more. Supernatants were collected and combined from both incubations, then centrifuged at 1200 rpm for 5 min at 4°C. RNA and protein were isolated from the pellet for further analysis. For culture purpose, the colons were cut into pieces and washed by DMEM for three times, then incubated with digestion buffer (250ug/ml collagenase type I and 500ug/ml Dispase II) for 1.5 h in 37°C. After the incubation, the cell suspension was passed through 100 um cell strainers (Corning). After washes, the cells were plated in dishes coated with rat tail tendon collagen type I overnight and were cultured in DMEM with 10% FBS and 1% penicillin/streptomycin. The next day, the cells were treated with NAC (5 mM) for subsequent experiments.

### Statistical Analysis

Data were analyzed using the student’s t-test and results are presented as the mean ± s.e.m (SEM) unless otherwise indicated. Pearson correlation coefficients were used to evaluate the relationships between SETD2 and gene expressions. Statistical analysis was performed using the GraphPad Prism software. A p value that was less than 0.05 was considered statistically significant for all data sets.

### Data Availability

All data are available from the authors upon reasonable request. RNA-Seq and ChIP-Seq raw data have been deposited in the Gene Expression Omnibus (GEO) under accession number GEO: GSE 151968.

## Results

### SETD2 Expression Becomes Decreased in IBD Patients and DSS Treated Mice

To explore a possible role of SETD2 in IBD, we analyzed SETD2 mRNA expression in public datasets of UC samples. The results indicate that the level of SETD2 mRNA was reduced in these samples compared with that in healthy controls (Figure 1A). To expand these observations, we performed real-time quantitative PCR (RT-qPCR) assays to determine SETD2 expression in colonic biopsies from IBD patients and healthy specimens. Consistent with the results obtained from dataset GSE9452, RT-qPCR analysis indicated that the SETD2 mRNA levels were significantly decreased in the IBD biopsies compared with their normal specimens (Figure 1B). To further evaluate the relevance of SETD2 in IBD, we measured SETD2 expression level in a colitis mouse model. DSS-induced colitis is a common mouse model that imitates the clinical pathology of IBD^40, 41^. We challenged wild-type mice with 3% DSS for 7 days and analyzed 2 days later (Figure 1C). The DSS-treated wild-type mice exhibited widespread damage and mild inflammation in the colon (Figure 1D). The excess inflammatory response observed in the DSS-treated wild-type mice was associated with a loss of mucus-producing goblet cells and antimicrobial peptide (AMP)-producing Paneth cells (Figure 1E). The two types of cells play an important role in intestinal antibacterial defense by releasing antimicrobial factors^42, 43^. Consistently, the DSS-treated colons produced significantly more proinflammatory cytokines and chemokines than the colons from the control littermates (Figure 1F). These results indicated that we successfully constructed a colitis mouse model. Subsequently, we evaluated SETD2 protein and RNA expressions in the intestine of colitis mice. Similar to what was seen in IBD patients, SETD2 was down-regulated in the intestine of DSS-induced colitis mice (Figure 1G-1H). Together, these findings suggest a causal link between SETD2 reduction and IBD pathogenesis.

**Figure 1.**
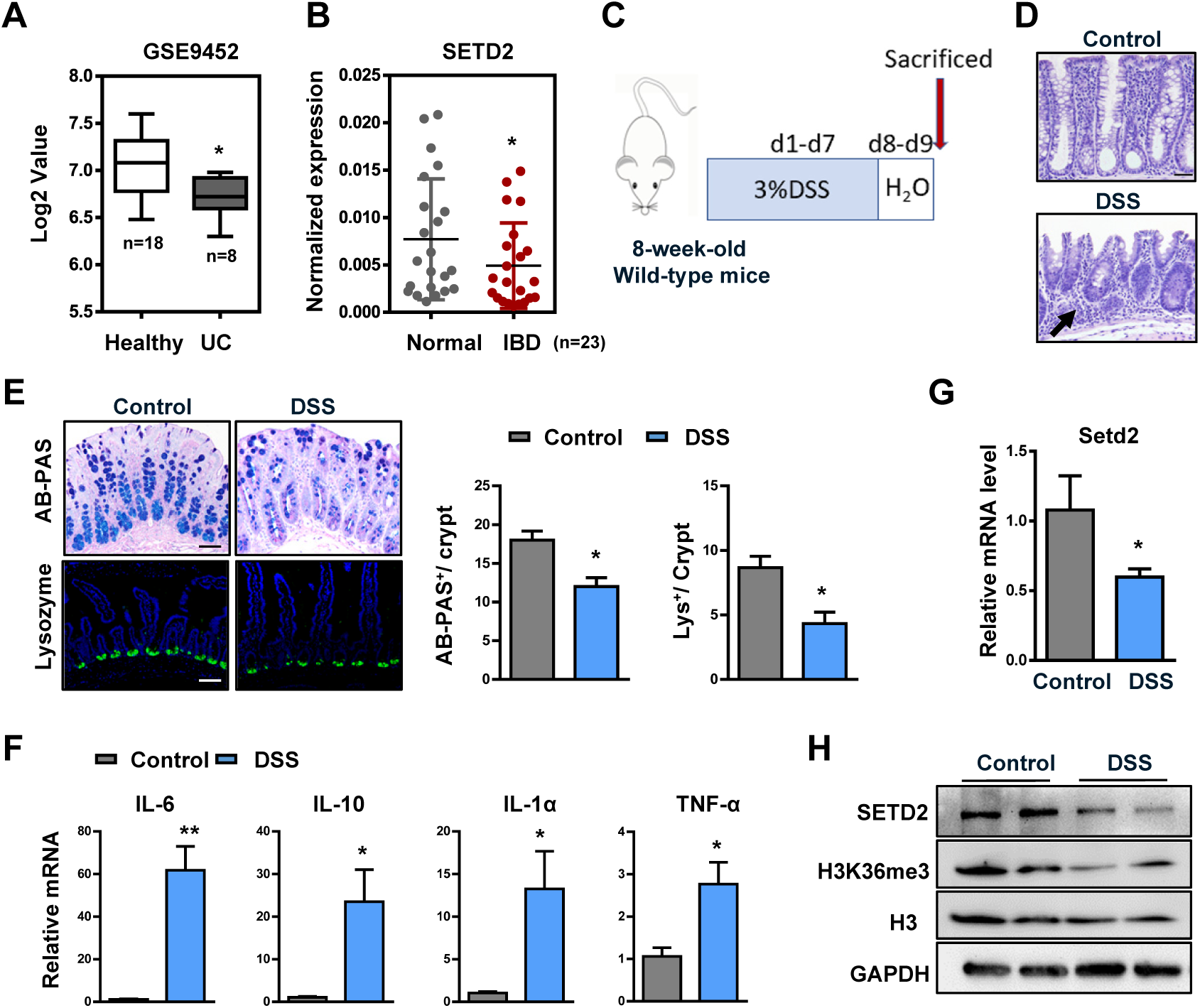
SETD2 expression is downregulated in IBD patients and DSS treated mice. **(A)** Boxplot of SETD2 expression levels in healthy controls and UC patients (using dataset GSE9452; n = 26). **(B)** RT-qPCR analysis of SETD2 transcripts in IBD specimens and healthy subjects (n = 23). **(C)** Schematic representation of the DSS protocol used to induce acute colitis. **(D-H)** Samples are all derived from 8-week-old wild-type mice with or without DSS treatment.**(D)** Representative images of H&E-stained colon sections as indicated. Scale Bars: 100um. **(E)** Alcian blue-Periodic acid Schiff (AB-PAS; goblet cells) staining and lysozyme (Lys; Paneth cells) staining in the colon as indicated, and quantitation results are shown in the right (n=3). Scale Bars: 100um. **(F)** RT-qPCR analysis of whole colon homogenates to assess cytokine and chemokine production (n = 4).**(G)** RT-qPCR analysis of SETD2 mRNA in control and DSS-treated wild-type mice(n=4). **(H)** Immunoblot analyses of SETD2 expression in control and DSS-treated wild-type mice are shown. The data represent the mean ± S.E.M, and statistical significance was determined by a two-tailed Student’s t-test. * p < 0.05.

### *Setd2*^*Vil-KO*^ >Mice Are More Susceptible to DSS-Induced Colitis

To assess a role of SETD2 in colonic inflammation, we crossed *Setd2-flox* mice (*Setd2*^*f/f*^ mice) with *Villin-Cre* mice to obtain an intestinal epithelium-specific *Setd2* knockout mouse strain (*Villin-Cre*; *Setd2 flox/flox* mice, hereinafter referred to as *Setd2*^*Vil-KO*^ mice). *Setd2*^*Vil-KO*^ mice were born with the correct Mendelian frequency. As expected, *Setd2* was efficiently ablated and H3K36me3 was substantially reduced in the intestinal epithelium of *Setd2*^*Vil-KO*^ mice by immunohistochemical staining and immunoblot validation (Figure 2A-2B). Histological examination of 8-week-old *Setd2*^*Vil-KO*^ mice revealed no obvious abnormalities in the colon crypt (Supplementary Figure 1A). However, as compared to the *Setd2*^*f/f*^ mice, the levels of several proinflammatory cytokines and chemokines (IL-1α, CXCL1) were significantly higher in the colonic sections of *Setd2*^*Vil-KO*^ mice (Supplementary Figure 1B). Meanwhile, *Setd2*^*Vil-KO*^ mice exhibited abnormal decreases of mucus-producing goblet cells and AMP-producing Paneth cells (Supplementary Figure 1C). These observations were further confirmed by RT-qPCR analysis (Supplementary Figure 1D), suggesting that *Setd2* deficiency in the intestine specifically causes a decrease in the number of secretory cells, which in turn leads to a disturbance in the homeostasis of the intestinal epithelium.

**Figure 2.**
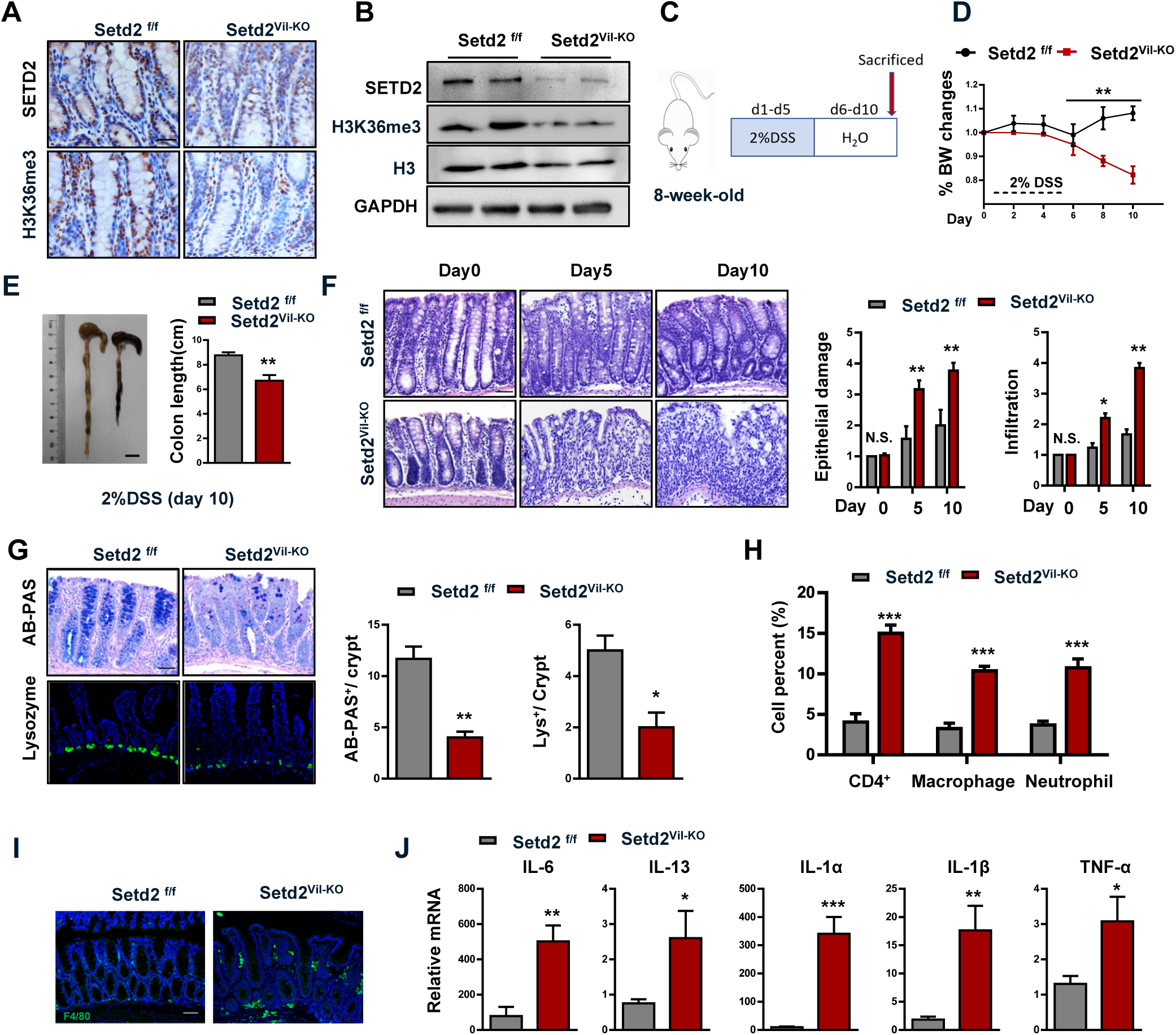
Loss of *Setd2* in IECs aggravates DSS-induced colitis in mice. **(A-B)** Immunohistochemical **(A)** and immunoblot analyses **(B)** of SETD2 and H3K36me3 expressions are shown. Scale Bars: 50um. **(C)** Schematic representation of the DSS protocol used to induce acute colitis. **(D, E)** *Setd2*^*Vil-KO*^ and *Setd2*^*f/f*^ mice were fed 2% DSS in drinking water and loss of body weights **(D)** and colon length **(E)** are recorded (n=5). Scale Bars: 1cm. **(F)** H&E stained sections of colon tissues collected on day 0, 5 and 10 from 2% DSS-treated mice, and the quantitation of DSS-induced epithelial damage and immune cell infiltration are shown in the right. Scale Bars: 100um. **(G)** Alcian blue-Periodic acid Schiff (AB-PAS; goblet cells) staining and lysozyme (Lys; Paneth cells) staining in the colon as indicated. Scale Bars: 100um. **(H)** After 5 d of DSS treatment, colonic lamina propria cells from *Setd2*^*Vil-KO*^ and *Setd2*^*f/f*^ mice are analyzed by flow cytometry for CD4 + T cells, CD11b +; F4/80 + macrophages, and CD11b + ; Gr-1 + neutrophils (n = 3). **(I)** Colon sections from mice treated with DSS were stained for F4/80 to detect macrophages. Scale Bars: 100um. **(J)** Relative mRNA expression levels of inflammatory mediators in whole colon of *Setd2*^*Vil-KO*^ and *Setd2*^*f/f*^ mice were determined by RT-qPCR (n=7). The data represent the mean ± S.E.M, and statistical significance was determined by a two-tailed Student’s t-test. * p < 0.05, ** p < 0.01 and *** p < 0.001. N.S., Not Significant.

To further define the role of SETD2 in colitis, we assessed the consequence of *Setd2* loss in acute colitis by challenging *Setd2*^*f/f*^ and *Setd2*^*Vil-KO*^ mice with 2% DSS and then monitored their susceptibility by histological and morphological characteristics (Figure 2C). After 5 days of DSS administration, *Setd2*^*Vil-KO*^ mice lost more body weight than *Setd2*^*f/f*^ mice (Figure 2D), suggesting that *Setd2*^*Vil-KO*^ mice probably had enhanced inflammation and intestinal damage, as body weight loss is one of the characteristics for the severity of DSS-induced colitis^44, 45^. In addition, macroscopic dissection revealed significantly shorter colons in the *Setd2*^*Vil-KO*^ mice compared with the *Setd2*^*f/f*^ mice (Figure 2E). On further histopathological examination, the *Setd2*^*Vil-KO*^ mice exhibited widespread damages in the colon, with more ulcerations as compared with *Setd2*^*f/f*^ mice. At both 5 and 10 days post DSS treatment, the *Setd2*^*f/f*^ colon showed minimal to mild inflammation, while colon from *Setd2*^*Vil-KO*^ mice displayed moderate to severe inflammation, with many areas of complete crypt loss and erosions (Figure 2F). The excess inflammatory response observed in the *Setd2*^*Vil-KO*^ mice was also associated with a loss of goblet cells and Paneth cells (Figure 2G). Since *Setd2*^*Vil-KO*^ mice showed higher epithelial damage and inflammation, we then analyzed immune cell infiltration by performing flow cytometry. At 5 days post DSS treatment, we observed a higher number of CD4 + T cells, neutrophils, and macrophages in *Setd2*^*Vil- KO*^ mice compared with control mice (Figure 2H). Notably, the immune cell infiltration also extended to the entire colon, as evidenced by the accumulation of F4/80-positive cells in relatively non-inflamed areas of the *Setd2*^*Vil-KO*^ colons (Figure 2I). It is well-known that infiltrating immune cells produce cytokines and chemokines to resolve the inflammation process^46, 47^. As expected, the DSS-treated *Setd2*^*Vil-KO*^ colons produced prominently more proinflammatory cytokines and chemokines than the colons from the *Setd2*^*f/f*^ littermates (Figure 2J). Collectively, these results demonstrate that depletion of *Setd2* leads to exacerbated colitis, suggesting that SETD2 plays a pivotal role in resolving inflammatory damages in the colonic epithelium.

### *Setd2* Deficiency Drives Inflammation-Associated CRC

The observation that *Setd2*^*Vil-KO*^ mice suffered from the sustained inflammation, prompted us to investigate a possible role of SETD2 in colitis-associated tumorigenesis. We induced CRC by injecting the DNA-methylating agent azoxymethane (AOM), followed by three cycles of 2% DSS treatments, and each cycle consisted of 5 d of 2%DSS water followed by 14 d of water alone. We recorded the changes in body weight throughout the duration of the DSS treatment, and determined the tumor burden after 90 days of AOM treatment. We found that *Setd2*^*Vil-KO*^ mice suffered from chronic inflammation and lost noticeably more body weight compared with the control mice (Figure 3A). Compared with the *Setd2*^*f/f*^ mice, *Setd2*-deficient mice had more macroscopic polypoid lesions, which were on average threefold larger in size (Figure 3B). Histological analysis revealed that around 50% of polyps in *Setd2*^*Vil-KO*^ mice were classified as high-grade dysplasia. However, the lesions developed in *Setd2*^*f/f*^ mice were mainly graded as low-grade dysplasia or hyperplasia (Figure 3C). Accordingly, the colons of *Setd2*^*Vil-KO*^ mice exhibited higher proliferative rates, indicated by Ki67 staining, than those in the control mice (Figure 3D). Thus, *Setd2* ablation strongly facilitates the development and progression of inflammation associated CRC.

**Figure 3.**
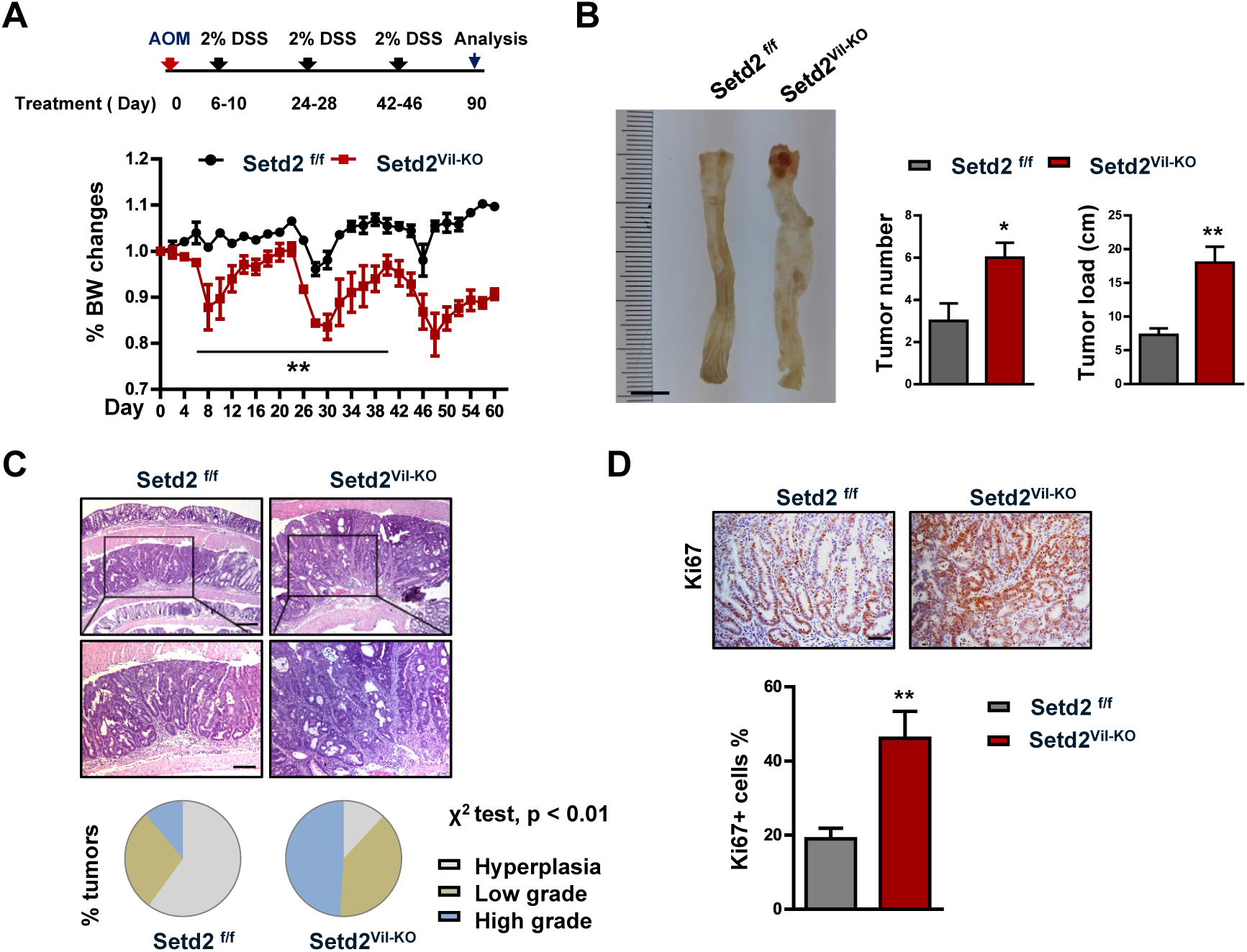
*Setd2* loss promotes the inflammation-associated CRC. **(A)** *Setd2*^*Vil-KO*^ and *Setd2*^*f/f*^ mice were injected with AOM on day 0 (2 months of age) and treated with 2% DSS during three 4-day cycles as indicated. Body weight changes are recorded as indicated (n = 4 per genotype). **(B)** At day 90 after the AOM injection, the mice were sacrificed to examine the tumor burden (n = 5 per genotype). Scale Bars: 1cm. **(C)** Representative images of colons and the overall grading of the tumors in each genotype (χ^2^ test) are shown (n = 5 per genotype). Scale Bars: upper, 200um; bottom, 100um. **(D)** Representative images of Ki67 staining in the tumors from AOM/DSS-treated *Setd2*^*Vil-KO*^ and *Setd2*^*f/f*^ mice are shown (n=5). Scale Bars: 50um. The data represent the mean ± S.E.M, and statistical significance was determined by a two-tailed Student’s t-test. * p < 0.05, ** p < 0.01.

### Disruption of *Setd2* Induces Apoptosis and Barrier Dysfunction in the Intestinal Epithelium

Given that intestinal epithelial barrier dysfunctions could frequently contribute to gut inflammation^6, 8, 9^, and that there were increased epithelial damages in the absence of *Setd2* (Figure 2F), we investigated whether the loss of *Setd2* compromised the barrier integrity by assessing the distribution of junction proteins and permeability of intestinal epithelial cells. We examined the distribution of the tight junction protein 1 (ZO-1) and E-cadherin in *Setd2*^*f/f*^ and *Setd2*^*Vil-KO*^ mice, two markers to reveal the integrity of junction structure and epithelial barriers. In contrast to the controls, the *Setd2*-deficient colons showed partially disrupted or discontinuous ZO-1 and E-cadherin staining (Figure 4A), indicating that an intact mucosal barrier was compromised in *Setd2*^*Vil-KO*^ mice. This observation was further strengthened by Western blot assays showing reduced expression of ZO-1, Claudin-1 and E-cadherin proteins in *Setd2*^*Vil-KO*^ mice compared with control mice (Figure 4B). In accordance with the severe ulceration (Figure 2F), intestinal permeability was markedly increased in *Setd2*^*Vil-KO*^ mice, evidenced by the enhanced fluorescence in the serum of DSS-treated *Setd2*^*Vil-KO*^ mice fed with FITC-labeled dextran (Figure 4C). Thus, these results indicate that *Setd2* silencing in colons leads to epithelial barrier dysfunction and colonic leakage.

**Figure 4.**
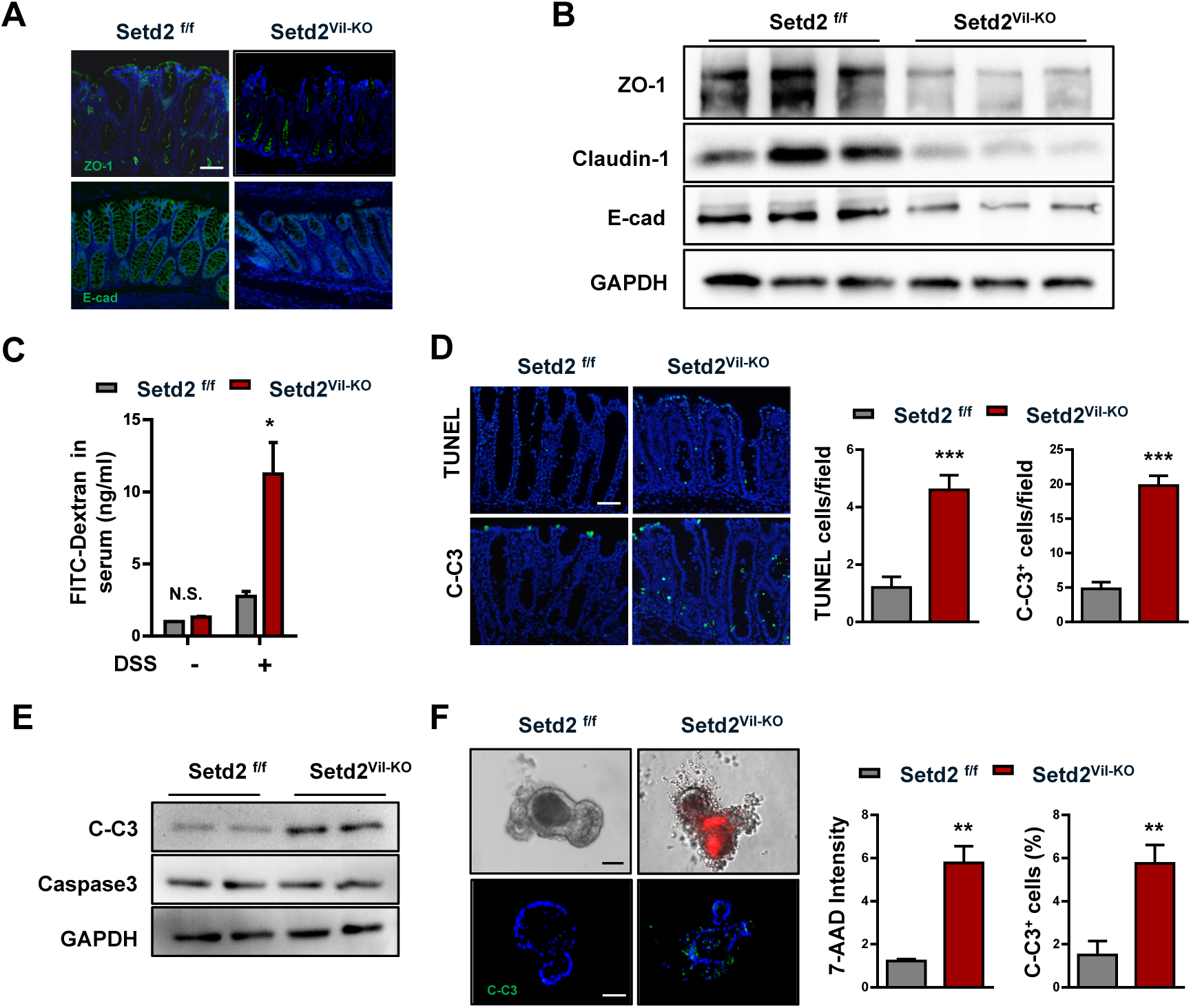
Loss of *Setd2* in IECs induces cell apoptosis and barrier dysfunction. **(A)** Representative ZO-1 and E-cadherin staining in colon sections from DSS-treated (5 d) *Setd2*^*Vil-KO*^ and *Setd2*^*f/f*^ mice. Scale Bars: 100um. **(B)** Immunoblotting analysis of the indicated proteins in IECs isolated from *Setd2*^*Vil-KO*^ and *Setd2*^*f/f*^ mice. **(C)** Colonic permeability was measured by the concentration of FITC-dextran in the blood serum (n=3). **(D)** TUNEL (Upper) and cleaved caspase 3 (Lower) staining of colon sections from DSS-treated (5 d) *Setd2*^*Vil-KO*^ and *Setd2*^*f/f*^ mice, and quantitation results are shown in the right (n=5). Scale Bars: 50um. **(E)** Western blot analysis of the indicated proteins in IECs isolated from DSS treated (5 d) *Setd2*^*Vil-KO*^ and *Setd2*^*f/f*^ mice. **(F)** Intestinal organoids derived from *Setd2*^*Vil-KO*^ and *Setd2*^*f/f*^ mice. After 5 d of differentiation, 7-AAD-stained organoids were imaged, and quantitations of the fluorescence density are shown in the right (n=4). Scale Bars: 20um. The data represent the mean ± S.E.M, and statistical significance was determined by a two-tailed Student’s t-test. * p < 0.05, ** p < 0.01, *** p < 0.001. N.S., Not Significant.

Considering that epithelial apoptosis is one of the mechanisms by which DSS can induce intestinal inflammation or colitis, and that the epithelial barrier is disrupted in the absence of *Setd2* (Figure 4C), we examined possible defects in epithelial cell survival in the *Setd2*^*Vil-KO*^ mice. On the histopathological examination, the numbers of TUNEL- and cleaved caspase-3-positive cells were significantly higher in the colon sections from the *Setd2*^*Vil-KO*^ mice than in those from the *Setd2*^*f/f*^ mice (Figure 4D). This observation was further strengthened by western blotting showing markedly elevated signaling intensities of cleaved caspase-3 in *Setd2*^*Vil-KO*^ mice compared with *Setd2*^*f/f*^ mice (Figure 4E). We next established intestinal organoid cultures to assess whether loss of *Setd2* induces apoptosis in a cell-intrinsic manner. 7-aminoactinomycin D (7- AAD) whole-mount staining and cleaved caspase-3 immunostaining displayed a substantial increase in apoptosis in *Setd2*-depleted cells relative to the controls (Figure 4F). These data emphasize that SETD2 is intrinsically required for the colonic epithelial homeostasis, and the loss of *Setd2* results in the barrier dysfunction and the subsequent inflammation.

### SETD2 Modulates ROS Homeostasis to Regulate Colon Inflammation

To further investigate the molecular mechanisms in which SETD2 regulates intestinal epithelial homeostasis, we performed RNA-seq analysis. Next-generation sequencing using RNA from *Setd2*^*Vil-KO*^ and *Setd2*^*f/f*^ IECs after 4 d of DSS treatment respectively revealed that the global transcriptome was changed dramatically in *Setd2*^*Vil-KO*^ IECs compared to the *Setd2*^*f/f*^ IECs, indicating a significant function of SETD2 in IECs (Figure 5A). Among a total of 18168 genes expressed, 644 genes were up-regulated, and 605 genes were down-regulated (Fold change > 1.25) in *Setd2*^*Vil-KO*^ IECs. Gene Ontology (GO) term analysis of the expression profile indicated that there was a significant enrichment of genes linked to apoptotic regulation, inflammatory response and oxidation activity (Figure 5A). These results were validated by RT-qPCR (Supplementary Figure 2A). Notably, the mRNA expression levels of some antioxidant genes were significantly down-regulated, suggesting increased oxidative stress in the absence of *Setd2*. To confirm this finding, we performed 8-oxo-2’-deoxyguanosine (8-OHdG) staining to examine oxidative stress in colon. We detected that 8-OHdG level was significantly increased in the colons of *Setd2*^*Vil-KO*^ mice after DSS treatment (Figure 5B). Oxidative stress is associated with intestinal inflammation and CRC pathogenesis, and ROS plays an important role in apoptosis induction and inflammation damage under both physiologic and pathologic conditions^12, 13^. To study the role of ROS in SETD2-mediated inflammation, we isolated the IECs from *Setd2*^*Vil-KO*^ and *Setd2*^*f/f*^ mice after DSS treatment. Flow cytometry analysis by staining 2′,7′-dichlorodihydrofluorescein diacetate (H2DCFDA, a fluorescent probe that reacts with ROS) indicated that *Setd2*-deficency IECs produced significantly more amount of ROS relative to that in the control IEC cells (Figure 5C). These results indicate that ROS is a necessary player in SETD2-mediated colitis.

**Figure 5.**
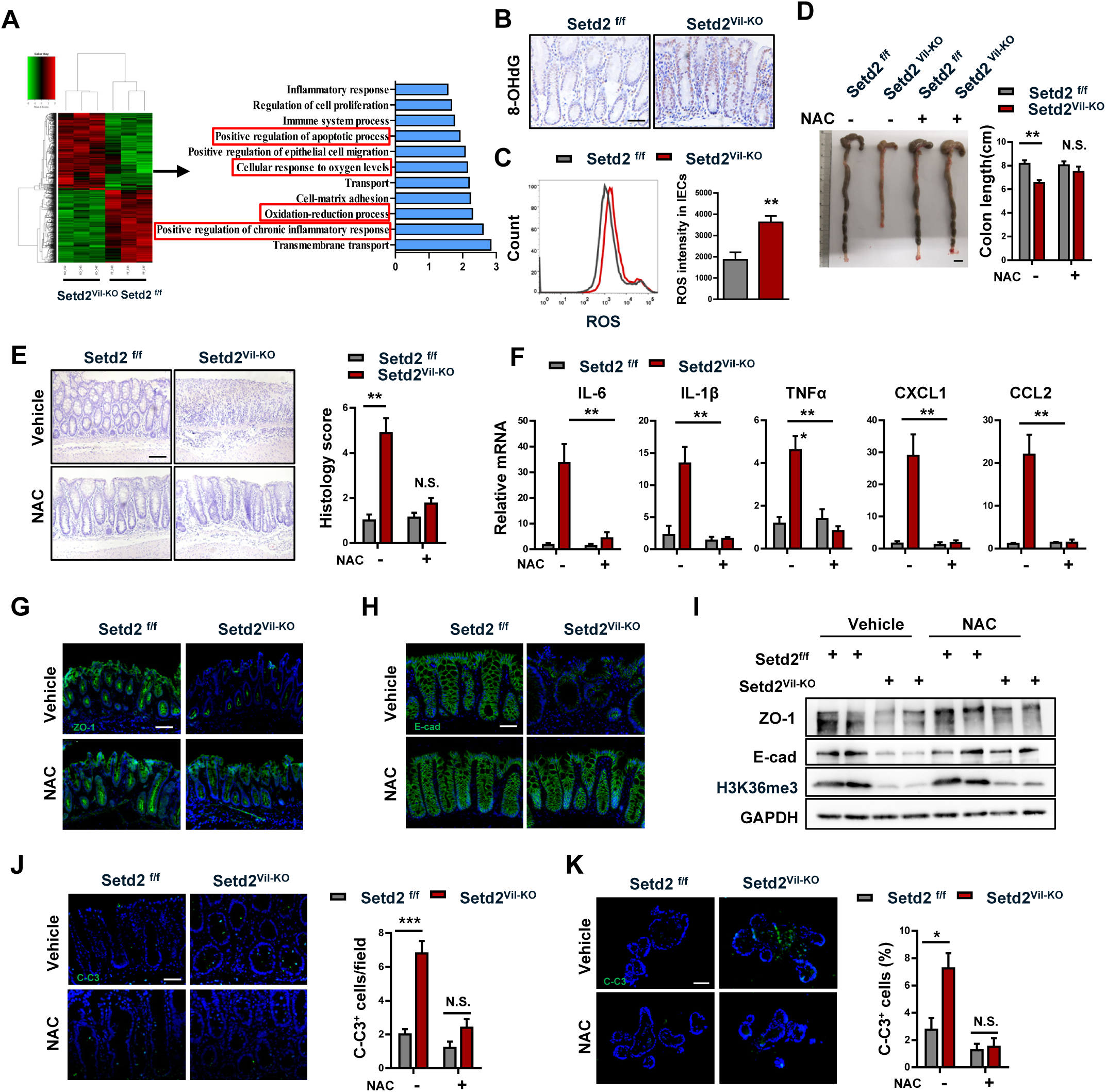
SETD2 mediates ROS reaction to control cell apoptosis and colon inflammation. **(A)** Heat map of RNA-seq data to compare the gene expression of colons from *Setd2*^*Vil- KO*^ and *Setd2*^*f/f*^ mice. Go term analysis of gene expression changes in the right. **(B)** 8-OHdG staining from DSS-treated *Setd2*^*Vil-KO*^ and *Setd2*^*f/f*^ mice. Scale Bars: 50um. **(C)** Histograms and quantifications of ROS in IECs isolated from DSS-treated *Setd2*^*Vil- KO*^ and *Setd2*^*f/f*^ mice. **(D-J)** 8-week-old *Setd2*^*Vil-KO*^ and *Setd2*^*f/f*^ mice were pretreated with NAC (100 mg/kg) every day from 4 days before the treatment with 2% DSS to the end of the experiment. Colon length (**D)**, H&E staining (**E)**, RT-qPCR analysis **(F)**, ZO-1 staining **(G)**, E-cadherin staining **(H)**, Western Blotting analysis **(I)** and cleaved caspase-3 staining **(J)** as indicated. Scale Bars: 100um (e, g, h); 50um (j). **(K)** Organoids were isolated from 8-week-old *Setd2*^*Vil-KO*^ and *Setd2*^*f/f*^ mice with or without NAC(5mM) treatment, and cleaved caspase-3 staining. Scale Bars: 20um. The data represent the mean ± S.E.M, and statistical significance was determined by a two-tailed Student’s t-test. * p < 0.05, ** p < 0.01, *** p < 0.001. N.S., Not Significant.

To provide evidence that ROS indeed contributes to the barrier defects and colitis, we performed rescue assays to determine whether blocking ROS could reverse the inflammatory phenotype caused by the *Setd2* loss. Two-month-old *Setd2*^*Vil-KO*^ and *Setd2*^*f/f*^ mice were pretreated with NAC (100 mg/kg) every day from 4 days before treated with 2% DSS to the end of the experiment, except for the 5 days during which DSS was administered^12, 48^. According to colon length and histopathological staining (Figure 5D-5E), the blockade of ROS production via the administration of NAC in the *Setd2*^*Vil-KO*^ mice completely eased the severity of inflammation, and the colonic morphology score was restored to the normal value. As judged by the expression levels of proinflammatory cytokines and chemokines, the severe inflammatory response occurring in the *Setd2*^*Vil-KO*^ mice was largely mitigated by the ROS inhibition (Figure 5F). Additionally, the NAC treatment restored the numbers of Paneth and goblet cells to levels similar to those in the control mice (Supplementary Figure 2B-2C). Besides, NAC-treated *Setd2*^*Vil-KO*^ mice exhibited a less barrier disruption and epithelial cell apoptosis than *Setd2*^*f/f*^ mice (Figure 5G-5J). In addition, the relieved epithelial apoptosis was also repeated by intestinal organoid culture experiment, indicated by cleaved caspase 3 staining (Figure 5K). Altogether, these results suggest that the blockade of ROS could reverse the phenotype of epithelial barrier disruption and colon inflammation in *Setd2*^*Vil-KO*^ mice. Thus, ROS plays an important role in SETD2-mediated colitis.

### SETD2-Mediated H3K36me3 Facilitates the Transcriptions of Antioxidant Genes to Maintain ROS Homeostasis

To elucidate the molecular basis by which SETD2 modulated ROS, we immunoprecipitated H3K36me3-bound chromatin in *Setd2*^*Vil-KO*^ and *Setd2*^*f/f*^ IECs after 4 d of DSS treatment and analyzed the precipitated DNA by deep sequencing. 178919 and 217927 H3K36me3 peaks were identified in *Setd2*^*Vil-KO*^ and *Setd2*^*f/f*^ IECs, respectively. The altered density of H3K36me3 intervals (*Setd2*^*Vil-KO*^ versus *Setd2*^*f/f*^) were widely localized within whole genomic regions, including gene bodies, promoters, and intergenic zones (Figure 6A-6B), indicating that SETD2 is responsible for the H3K36 trimethylaiton on these regions. To correlate the chromatin binding with the transcriptional regulation, we integrated the ChIP-seq data with the expression profile and the Venn diagrams indicated that 906 (including downregulated and upregulated) genes showed direct H3K36me3 occupancies and expression changes upon the *Setd2* ablation (Figure 6C). GO term analysis revealed that these overlapping 906 genes were still closely related to oxidative stress pathway (Figure 6C). As shown in figure 5, ablation of *Setd2* increased ROS production and down-regulated the expression levels of antioxidant genes, suggesting that Setd2/H3K36me3 might positively regulate some antioxidant-coding genes related to ROS production. As expected, antioxidant genes including *Prdx3, Prdx6, Gclm*, and *Srxn1* were listed among the 906 genes described above, and their downregulation was validated in colon epithelial cells collected from *Setd2*^*Vil-KO*^ mice, comparing with cells from *Setd2*^*f/f*^ mice (Figure 6D). Direct H3K36me3 occupancies within these candidate gene loci were also seen in Genome Browser tracks (Figure 6E). We further validated the existence of H3K36me3 bindings at the gene bodies of *Prdx3, Prdx6, Gclm* and *Srxn1* by the ChIP-qPCR assays, and found that the intensity of H3K36me3 bindings within these gene loci decreased along with the loss of *Setd2* (Figure 6F). Consistent with these results, there were significant positive correlations between mRNA levels of SETD2 and that of PRDX3, PRDX6, GCLM and SRXN1 respectively based on GSE57945 database (Figure 6G). Thus, *Setd2* loss could repress the transcription of antioxidant genes, which were important for antioxidant defenses. The peroxiredoxin [PRDX] family contains the most of important antioxidant proteins. PRDX6, a bifunctional 25-kDa protein, is one member of PRDXs^49^. Next, we sought to investigate whether restoration of PRDX6 in *Setd2*-depleted IECs would attenuate the overproduction of ROS and improve the cell survival as compared with *Setd2*-deleted IECs. As expected, overexpression of PRDX6 in *Setd2*-depleted IECs could significantly reduce the ROS production and the cleaved caspase-3 signals (Figure 6H-6I). Thus, ectopic *Prdx6* could alleviate the overproduction of ROS and improve the cellular survival in the *Setd2*-deficient intestinal epithelium. Together, these results indicate that SETD2 prevents colon inflammation and epithelial barrier defects via the regulation of antioxidant to restrain ROS over reaction.

**Figure 6.**
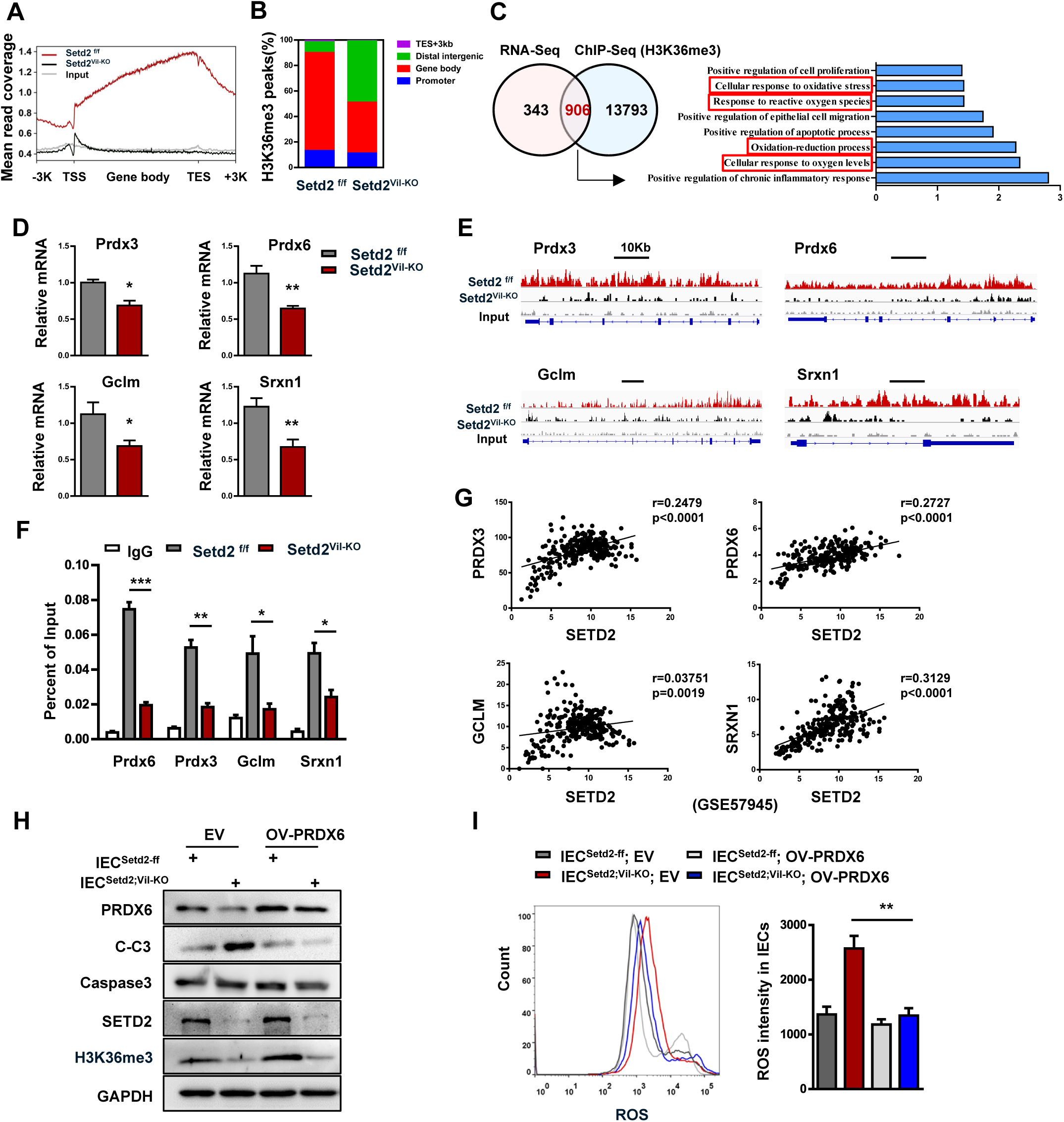
SETD2 regulates antioxidant protein via H3K36me3. **(A)** Normalized read density of H3K36me3 ChIP-seq signals of colons from *Setd2*^*Vil- KO*^ and *Setd2*^*f/f*^ mice from 3 kb upstream of the TSS to 3 kb downstream of the TES. **(B)** Analysis of the occupancy of H3K36me3 ChIP-seq peaks in gene bodies and intergenic regions. **(C)** Venn diagram showing the number of genes harboring H3K36me3 binding and displaying expression changes in *Setd2*^*Vil-KO*^ IECs. Right panel shows the Go term analysis of the overlapping genes. **(D)** RT-qPCR analysis of antioxidant genes expression in IECs as indicated. **(E)** Snapshot of H3K36me3 ChIP-Seq signals at the *Prdx3, Prdx6, Gclm* and *Srxn1* gene loci in IECs isolated from DSS-treated (4 d) *Setd2*^*Vil-KO*^ and *Setd2*^*f/f*^ mice. **(F)** ChIP-qPCR analysis of H3K36me3 binding for *Prdx3, Prdx6, Gclm* and *Srxn1* loci in IECs from DSS-treated (4 d) *Setd2*^*Vil-KO*^ and *Setd2*^*f/f*^ mice, and IgG was used as the control. **(G)** Correlation between SETD2 and PRDX3, PRDX6, GCLM and SRXN1 expression levels in IBD specimens (GSE 57945). Statistical significance was determined using the Pearson correlation coefficient. **(H)** Western blot analysis of the indicated proteins in *IEC*^*Setd2-ff*^ and *IEC*^*Setd2;VIL-KO*^ with or without PRDX6 overexpression. **(I)** Histograms and quantifications of ROS in *IEC*^*Setd2-ff*^ and *IEC*^*Setd2;VIL-KO*^ with or without PRDX6 overexpression. The data represent the mean ± S.E.M, and statistical significance was determined by a two-tailed Student’s t-test. * p < 0.05, ** p < 0.01, *** p < 0.001.

Gut microbes also play an important role in driving colonic inflammatory responses^5, 50^. To rule out the possibility that the intestinal microbiota contributes to epithelial barrier damage caused by *Setd2* deficiency, we generated gut microbiota-depleted mice by treating *Setd2*^*Vil-KO*^ and *Setd2*^*f/f*^ mice with drinking water containing an antibiotic cocktail for 4 weeks and then treated them with 2%DSS^51, 52^. As compared with the *Setd2*^*f/f*^ mice, there was less bacterial richness but unchanged gut microbiota composition in untreated *Setd2*^*Vil-KO*^ mice (Supplementary Figure 3A-B). Interestingly, depletion of the gut microbiota by antibiotics eased the severity of colitis and reduced expression of proinflammatory cytokines and chemokines of *Setd2*^*Vil-KO*^ mice comparable to those observed in the *Setd2*^*f/f*^ mice (Supplementary Figure 3C-D). However, antibiotic-treated *Setd2*^*Vil-KO*^ mice still exhibited more severe barrier disruption and epithelial cell apoptosis than *Setd2*^*f/f*^ mice (Supplementary Figure 3E-J). These results suggested that depletion of the gut microbiota did not facilitate the recovery of intestinal barrier in *Setd2*^*Vil-KO*^ mice. Thus, the gut microbiota does not contribute to intestinal barrier disruption caused by *Setd2* deficiency. Considered together, SETD2 protects the colon epithelial barrier from inflammatory insults through a mechanism dependent on oxidative stress (Figure 7).

**Figure 7.**
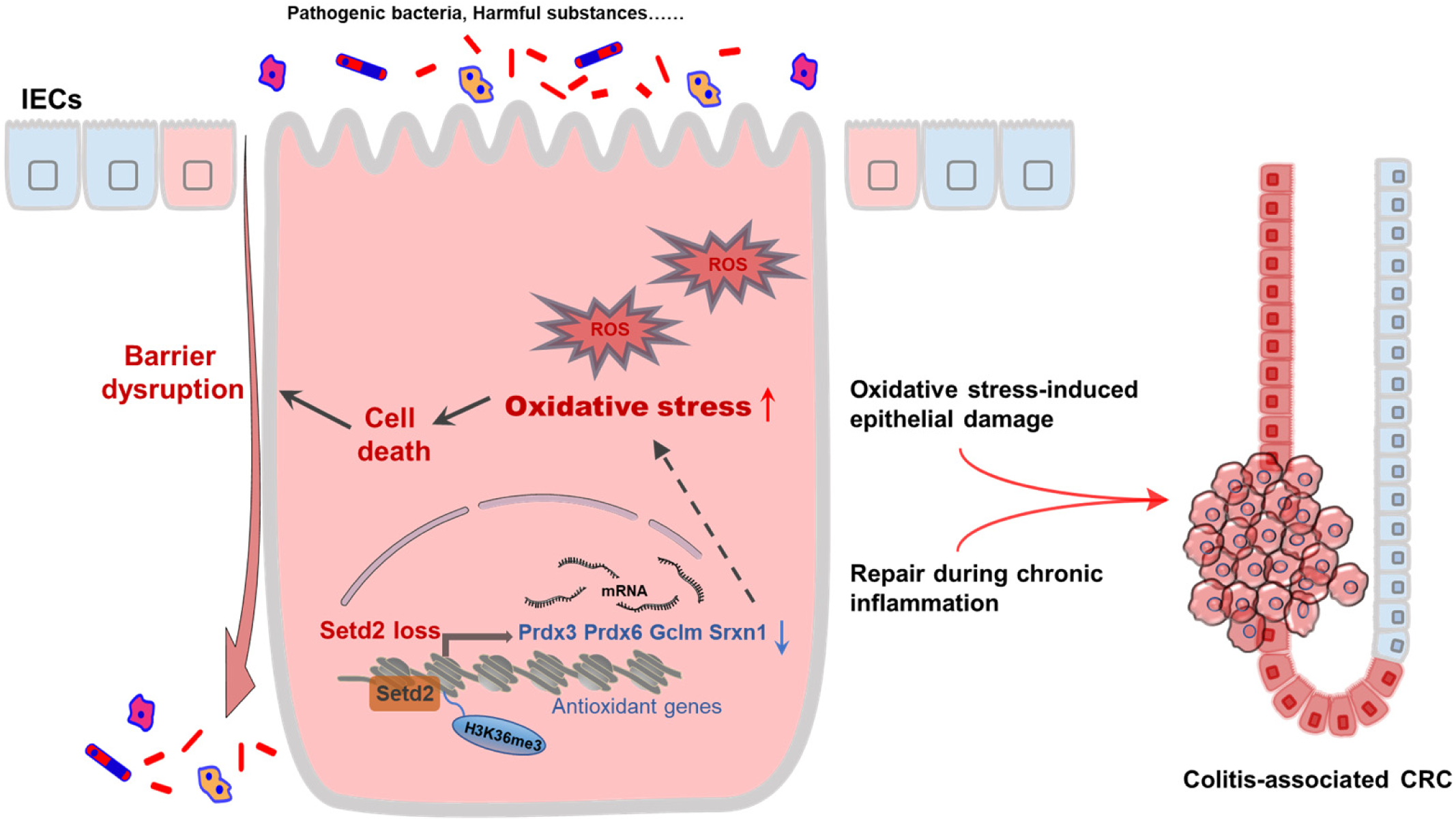
Loss of *Setd2* promotes oxidative stress-induced colonic inflammation and tumorigenesis. SETD2 directly controls multifaceted antioxidant genes including *Prdx3, Prdx6, Gclm* and *Srxn1* in intestinal epithelial cells (IECs). Thus, *Setd2* loss in IECs leads to insufficient antioxidant, which results in excess reactive oxygen species (ROS), and thereby compromises barrier integrity and promotes inflammation. Compensatory regeneration coupled with oxidative stress-induced epithelial damage promotes the malignant progression of CRC.

## Discussion

Although epigenetic dysregulation is recently believed to contribute to development of intestinal inflammation^24, 25, 53^ and epithelial barrier disruption is one of the major causes of IBD, whether epigenetic elements are drivers during the regulation of epithelial barrier integrity and colonic inflammation and tumorigenesis has not been genetically determined. In this study, we established mice with epithelium-specific deletion of *Setd2* (*Setd2*^*Vil-KO*^ mice), and then challenged with various inducing agents such as DSS and AOM to generate different types of models of colonic inflammation and tumorigenesis. We demonstrated clearly that deletion of *Setd2* that is responsible for one of the major histone modifications, disturbs intestinal epithelial homeostasis and worsens barrier defects by modulating oxidative stress, which promotes colonic inflammatory response and subsequently tumorigenesis (Figure 7). Moreover, the excess inflammatory damage occurring in the Setd2-depleted mice was largely alleviated by the ROS inhibition through antioxidant NAC treatment. Our results highlight the importance that SETD2 functions as a homeostatic regulator that integrates the intact epithelial barrier by modulating oxidative stress in the colon. Support for our findings comes from a recent report indicating that mutations of the *Setd2* gene occur in up to 17% in UC samples with a high risk of developing colorectal carcinoma ^28^. Thus, our study provides the first direct causal evidence that SETD2 plays a pivotal role in IBD and colitis-associated CRC.

The present study reveals a mechanism by which SETD2 inhibits ROS overreaction to maintain intestinal epithelial homeostasis against intestinal inflammation and tumorigenesis. Ulcerative colitis is usually characterized by mucosal barrier dysfunction, and intestinal barrier integrity is essential for preventing microbial-driven inflammatory response. Our study found that *Setd2* loss results in epithelial barrier dysfunction and colonic leakage, which promotes the subsequent inflammation response. ROS plays an important role in cell apoptosis induction and inflammation under both physiologic and pathologic conditions^12, 13, 19^. Mammalian cells are always exposed to ROS produced by environmental oxidants or endogenous metabolism^54, 55^. We found that *Setd2* deficiency in the colon epithelium leads to ROS accumulations, and then enhances epithelial cell death and barrier disruption. And this finding is further supported by the experiment of antioxidant NAC treatment. The NAC treatment in *Setd2*^*Vil-KO*^ mice attenuates the ROS reactions and relieves the inflammation symptoms. Therefore, excess ROS accumulation is mainly responsible for the colon inflammation observed in the *Setd2*^*Vil-KO*^ mice.

The notion that SETD2 regulates colon inflammation is via modulation of ROS homeostasis is further supported by our more detailed analysis that SETD2 facilitates the transcriptions of antioxidant genes. It is well known that SETD2 exerts epigenetic regulation via promoting the transcriptions of its downstream genes by marking gene bodies with H3K36me3 and facilitating the elongation of their pre-mRNA^29, 56^. At molecular level, we show that *Setd2* deficiency down-regulates the mRNA levels of the members of antioxidant genes including *Prdx3, Prdx6, Gclm* and *Srxn1* in the analysis of sequencing results. The current findings demonstrate that SETD2 attenuates colonic inflammation through maintaining the expression of antioxidant genes to suppress oxidative stress. Our work is further corroborated by the rescue experiment showing that overexpression of PRDX6 in *Setd2*-depleted IECs could reduce the ROS production and improve epithelial cell survival. Clinically, we also found that expression level of SETD2 in IBD samples is positively correlated with the expressions of PRDX3, PRDX6, GCLM and SRXN1, respectively. Therefore, our findings reveal for the first time the important role of SETD2-mediated H3K36me3 modification involved in the oxidative stress pathway in the maintenance of intestinal epithelial homeostasis, and provide insights into the molecular mechanisms underlying the intestinal inflammation and colitis-associated CRC with epigenetic disorders.

It is worth mentioning that SETD2 appears to interact with various proteins to influence transcription. Notably, our previous work reported that SETD2-mediated H3K36me3 could participate in cross-talks with other chromatin markers in oocytes, including H3K4me3 and H3K27me3 in transcribing regions^38^. In the present study, we demonstrate that expression of some antioxidant genes is regulated by SETD2-mediated H3K36me3. However, little is known about the epigenetic functions of other chromatin markers on regulating these genes in IBD. It would be interesting to study the physiological role of the cross-talk between H3K36me3 and other chromatin markers in IBD and colitis-associated CRC in the future. Moreover, recent studies have also revealed that SETD2 not only exerts its classical role in controlling gene transcription by mediating modifications of histone proteins, but also has the capacity to regulate cellular signaling through modification of non-histone substrates. For example, it has been reported that SETD2 could tri-methylate α-tubulin on K40 and mediate K525me1 of STAT1, which implicates a vital role in mitosis and antiviral immunity respectively^57, 58^. In the present study, we show that SETD2-mediated H3K36me3 promotes the expression of some antioxidant genes. However, it remains to be determined whether SETD2 can interact with these proteins and methylate them to affect the progression of IBD and tumorigenesis.

In summary, our findings highlight that SETD2 plays a critical role in the intestinal homeostasis and regulates barrier function and colon inflammation by modulating oxidative stress in the colon. We also established a SETD2/H3K36me3-deficient IBD mouse model that could be used for pre-clinical researches on colitis with epigenetic disorders. Given that SETD2 mutation accounts for 17% IBD patients with a high risk of developing colorectal carcinoma, our results may provide insight into our understanding of pharmaceutical investigation of these diseases associated with SETD2/H3K36me3 mutation.

## Abbreviations

AMP: antimicrobial peptide
AOM: azoxymethane
7-AAD: 7- aminoactinomycin D
CD: Crohn’s disease
CRC: colorectal cancer
DSS: dextran sodium sulfate
GO: gene ontology
H2DCFDA: 2′,7′-dichlorodihydrofluorescein diacetate
H3K36: histone H3 lysine 36
IBD: inflammatory bowel disease
IEC: intestinal epithelial cell
NAC: N-acetyl-l-cysteine
8-OHdG: 8-oxo-2’- deoxyguanosine
PRDX: peroxiredoxin
ROS: reactive oxygen species
RT-qPCR: Real-Time quantitative PCR
UC: ulcerative colitis
ZO-1: tight junction protein 1.

## Disclosures

All authors have declared that no conflict of interest exists.

## Acknowledgements

This study was supported by funds from Ministry of Science and Technology of the People’s Republic of China (2017YFA0102900 to W.Q.G.), National Natural Science Foundation of China (81772938 to L.L., 81872406 and 81630073 to W.Q.G.), State Key Laboratory of Oncogenes and Related Genes (KF01801 to L.L.), Science and Technology Commission of Shanghai Municipality (18140902700 and 19140905500 to L.L., 16JC1405700 to W.Q.G.), KC Wong foundation (to W.Q.G.) and Innovation Research Plan from Shanghai Municipal Education Commission (ZXGF082101 to L.L.). The study is also supported by Bio-ID Center, School of Biomedical Engineering, Shanghai Jiao Tong University.

## Notes

### Competing Interest Statement

The authors have declared no competing interest.

## References

1. Maloy KJ, Powrie F. Intestinal homeostasis and its breakdown in inflammatory bowel disease. Nature 2011;474:298–306.

2. Kaser A, Zeissig S, Blumberg RS. Inflammatory bowel disease. Annu Rev Immunol 2010;28:573–621.

3. Grivennikov SI, Greten FR, Karin M. Immunity, inflammation, and cancer. Cell 2010;140:883–99.

4. Gillen CD, Walmsley RS, Prior P, et al. Ulcerative colitis and Crohn’s disease a comparison of the colorectal cancer risk in extensive colitis. Gut 1994;35:1590–1592.

5. Xavier RJ, Podolsky DK. Unravelling the pathogenesis of inflammatory bowel disease. Nature 2007;448:427–34.

6. Rescigno M. The intestinal epithelial barrier in the control of homeostasis and immunity. Trends Immunol 2011;32:256–64.

7. Nowarski R, Jackson R, Gagliani N, et al. Epithelial IL-18 Equilibrium Controls Barrier Function in Colitis. Cell 2015;163:1444–56.

8. Schmitz H, Barmeyer C, Fromm M, et al. Altered Tight Junction Structure Contributes to the Impaired Epithelial Barrier Function in Ulcerative Colitis. Gastroenterology 1999;116:301–309.

9. Westbrook AM, Szakmary A, Schiestl RH. Mechanisms of intestinal inflammation and development of associated cancers: lessons learned from mouse models. Mutat Res 2010;705:40–59.

10. Sedelnikova OA, Redon CE, Dickey JS, et al. Role of oxidatively induced DNA lesions in human pathogenesis. Mutat Res 2010;704:152–9.

11. Moura FA, de Andrade KQ, Dos Santos JCF, et al. Antioxidant therapy for treatment of inflammatory bowel disease: Does it work? Redox Biol 2015;6:617–639.

12. Ravindran R, Loebbermann J, Nakaya HI, et al. The amino acid sensor GCN2 controls gut inflammation by inhibiting inflammasome activation. Nature 2016;531:523–527.

13. Barrett CW, Reddy VK, Short SP, et al. Selenoprotein P influences colitis-induced tumorigenesis by mediating stemness and oxidative damage. J Clin Invest 2015;125:2646–60.

14. Federico A, Morgillo F, Tuccillo C, et al. Chronic inflammation and oxidative stress in human carcinogenesis. Int J Cancer 2007;121:2381–6.

15. Tian T, Wang Z, Zhang J. Pathomechanisms of Oxidative Stress in Inflammatory Bowel Disease and Potential Antioxidant Therapies. Oxid Med Cell Longev 2017;2017:4535194.

16. Darnaud M, Dos Santos A, Gonzalez P, et al. Enteric Delivery of Regenerating Family Member 3 alpha Alters the Intestinal Microbiota and Controls Inflammation in Mice With Colitis. Gastroenterology 2018;154:1009–1023 e14.

17. Kinchen J, Chen HH, Parikh K, et al. Structural Remodeling of the Human Colonic Mesenchyme in Inflammatory Bowel Disease. Cell 2018;175:372–386 e17.

18. Pei R, Liu J, Martin DA, et al. Aronia Berry Supplementation Mitigates Inflammation in T Cell Transfer-Induced Colitis by Decreasing Oxidative Stress. Nutrients 2019;11.

19. Liu M, Sun T, Li N, et al. BRG1 attenuates colonic inflammation and tumorigenesis through autophagy-dependent oxidative stress sequestration. Nat Commun 2019;10:4614.

20. Okuda M, Li K, Beard MR, et al. Mitochondrial injury, oxidative stress, and antioxidant gene expression are induced by hepatitis C virus core protein. Gastroenterology 2002;122:366–75.

21. Kruidenier L, Kuiper I, Lamers CB, et al. Intestinal oxidative damage in inflammatory bowel disease: semi-quantification, localization, and association with mucosal antioxidants. J Pathol 2003;201:28–36.

22. Alenghat T, Osborne LC, Saenz SA, et al. Histone deacetylase 3 coordinates commensal-bacteria-dependent intestinal homeostasis. Nature 2013;504:153–7.

23. Theodoris CV, Li M, White MP, et al. Human disease modeling reveals integrated transcriptional and epigenetic mechanisms of NOTCH1 haploinsufficiency. Cell 2015;160:1072–86.

24. Epigenetics of inflammatory bowel disease. Gut 2000;47:302–306.

25. Hasler R, Feng Z, Backdahl L, et al. A functional methylome map of ulcerative colitis. Genome Res 2012;22:2130–7.

26. Cooke J, Zhang H, Greger L, et al. Mucosal genome-wide methylation changes in inflammatory bowel disease. Inflamm Bowel Dis 2012;18:2128–37.

27. Yongfeng Liu, Peng J, Tongyu Suna, Ni Li, et al. Epithelial EZH2 serves as an epigenetic determinant in experimental colitis by inhibiting TNFα-mediated inflammation and apoptosis. PNAS 2017;10:3796–3895.

28. Chakrabarty S, Varghese VK, Sahu P, et al. Targeted sequencing-based analyses of candidate gene variants in ulcerative colitis-associated colorectal neoplasia. Br J Cancer 2017;117:136–143.

29. Fontebasso AM, Schwartzentruber J, Zakrzewska M, et al. Mutations in SetD2 and genes affecting histone H3K36 methylation target hemispheric high-grade gliomas. Acta Neuropathol 2013;125:659–669.

30. Kanu N, Gronroos E, Martinez P, et al. SETD2 loss-of-function promotes renal cancer branched evolution through replication stress and impaired DNA repair. Oncogene 2015;34:5699–708.

31. Parker H, Rose-Zerilli MJ, Larrayoz M, et al. Genomic disruption of the histone methyltransferase SETD2 in chronic lymphocytic leukaemia. Leukemia 2016;30:2179–2186.

32. Mar BG, Bullinger LB, McLean KM, et al. Mutations in epigenetic regulators including SETD2 are gained during relapse in paediatric acute lymphoblastic leukaemia. Nat Commun 2014;5:3469.

33. Walter DM, Venancio OS, Buza EL, et al. Systematic In Vivo Inactivation of Chromatin-Regulating Enzymes Identifies Setd2 as a Potent Tumor Suppressor in Lung Adenocarcinoma. Cancer Res 2017;77:1719–1729.

34. Lee JJ, Park S, Park H, et al. Tracing Oncogene Rearrangements in the Mutational History of Lung Adenocarcinoma. Cell 2019;177:1842–1857 e21.

35. Wang L, Niu N, Li L, et al. H3K36 trimethylation mediated by SETD2 regulates the fate of bone marrow mesenchymal stem cells. PLoS Biol 2018;16:e2006522.

36. Ji Z, Sheng Y, Miao J, et al. The histone methyltransferase Setd2 is indispensable for V(D)J recombination. Nat Commun 2019;10:3353.

37. niu n, lu P, Yang Y, et al. Loss of Setd2 promotes Kras-induced acinar-to-ductal metaplasia and epithelia–mesenchymal transition during pancreatic carcinogenesis Gut 2019;0:1–12.

38. Xu Q, Xiang Y, Wang Q, et al. SETD2 regulates the maternal epigenome, genomic imprinting and embryonic development. Nat Genet 2019;51:844–856.

39. Zuo X, Rong B, Li L, et al. The histone methyltransferase SETD2 is required for expression of acrosin-binding protein 1 and protamines and essential for spermiogenesis in mice. J Biol Chem 2018;293:9188–9197.

40. Wirtz S, Neufert C, Weigmann B, et al. Chemically induced mouse models of intestinal inflammation. Nat Protoc 2007;2:541–6.

41. Wirtz S, Neurath MF. Mouse models of inflammatory bowel disease. Adv Drug Deliv Rev 2007;59:1073–83.

42. Wlodarska M, Thaiss CA, Nowarski R, et al. NLRP6 inflammasome orchestrates the colonic host-microbial interface by regulating goblet cell mucus secretion. Cell 2014;156:1045–59.

43. Bevins CL, Salzman NH. Paneth cells, antimicrobial peptides and maintenance of intestinal homeostasis. Nat Rev Microbiol 2011;9:356–68.

44. Okayasu I, Hatakeyama S, Yamada M, et al. A novel method in the induction of reliable experimental acute and chronic ulcerative colitis in mice. Gastroenterology 1990;98:694–702.

45. Gupta J, del Barco Barrantes I, Igea A, et al. Dual function of p38alpha MAPK in colon cancer: suppression of colitis-associated tumor initiation but requirement for cancer cell survival. Cancer Cell 2014;25:484–500.

46. Ramirez-Carrozzi V, Sambandam A, Luis E, et al. IL-17C regulates the innate immune function of epithelial cells in an autocrine manner. Nat Immunol 2011;12:1159–66.

47. Steinke JW, Borish L. 3. Cytokines and chemokines. J Allergy Clin Immunol 2006;117:S441–5.

48. Cha H, Lee S, Hwan Kim S, et al. Increased susceptibility of IDH2-deficient mice to dextran sodium sulfate-induced colitis. Redox Biol 2017;13:32–38.

49. Patel P, Chatterjee S. Peroxiredoxin6 in Endothelial Signaling. Antioxidants (Basel) 2019;8.

50. Belkaid Y, Hand TW. Role of the microbiota in immunity and inflammation. Cell 2014;157:121–41.

51. Rakoff-Nahoum S, Paglino J, Eslami-Varzaneh F, et al. Recognition of commensal microflora by toll-like receptors is required for intestinal homeostasis. Cell 2004;118:229–41.

52. Morgun A, Dzutsev A, Dong X, et al. Uncovering effects of antibiotics on the host and microbiota using transkingdom gene networks. Gut 2015;64:1732–43.

53. Ray G, Longworth MS. Epigenetics, DNA Organization, and Inflammatory Bowel Disease. Inflamm Bowel Dis 2019;25:235–247.

54. Shadel GS, Horvath TL. Mitochondrial ROS signaling in organismal homeostasis. Cell 2015;163:560–9.

55. Noubade R, Wong K, Ota N, et al. NRROS negatively regulates reactive oxygen species during host defence and autoimmunity. Nature 2014;509:235–9.

56. Neri F, Rapelli S, Krepelova A, et al. Intragenic DNA methylation prevents spurious transcription initiation. Nature 2017;543:72–77.

57. Park IY, Powell RT, Tripathi DN, et al. Dual Chromatin and Cytoskeletal Remodeling by SETD2. Cell 2016;166:950–962.

58. Chen K, Liu J, Liu S, et al. Methyltransferase SETD2-Mediated Methylation of STAT1 Is Critical for Interferon Antiviral Activity. Cell 2017;170:492–506 e14.

