## supplemental information for "The Histone Methyltransferase SETD2 Modulates Oxidative Stress to Attenuate Colonic Inflammation and Tumorigenesis in Mice"

#### Supplemental Methods

##### *RNA Isolation and Real-Time PCR Analysis*

Total RNA was isolated with RNeasy Mini Kit (Qiagen) according to the manufacturer's instructions, and was reverse transcribed into cDNA with the PrimeScript™ RT Reagent Kit (TakaRa). SYBR Green Master Mix reagents (TakaRa) were used for the real-time PCR. The primers used are listed in Supplementary Table 1.

##### *Western Blotting*

Cells or tissues were lysed with a RIPA buffer supplemented with protease and phosphatase inhibitors. Equal amount of protein samples were separated by 8% SDS-polyacrylamide gel electrophoresis (PAGE). The gel was transferred to a NC membrane and then incubated with primary antibodies overnight, followed by an appropriate secondary antibody. Chemiluminescent signal was measured using the Super Signal West Pico substrate (Millipore). The antibodies used are listed in Supplementary Table 2.

##### *Histology and Immunohistochemistry Staining*

Colons were dissected and flushed with PBS solution to remove faecal contents. For paraffin embedding, tissue segments were fixed in 4% paraformaldehyde overnight at 4℃ before paraffin embedding according to standard methods. For frozen section, tissue segments were embedded in OCT after 4% paraformaldehyde fixation and 30% sucrose solution sedimentation. In immunochemistry experiments, 5μm sections were prepared and dewaxed in xylene, rehydrated, and boiled for 15minutes in sodium citric buffer (pH6.0). Endogenous peroxidase activity was blocked in 3% hydrogen peroxide and then incubated in a 2% BSA/PBS solution before overnight incubation in primary antibody. After three times of PBS wash, sections were incubated in horseradish peroxidase-conjugated secondary antibodies for 1 hour, developed via DAB, (GeneTech Inc., Shanghai, China). The preparations were then counterstained with hematoxylin, dehydrated and finally mounted in Cytoseal. The antibodies used are listed in Supplementary Table 2.

***ChIP-qPCR Assays***

IECs were isolated from *Setd2*$^{f/f}$ mice and *Setd2*$^{Vil\text{-}KO}$ mice as described above. IECs were cross-linked with 1% formaldehyde for 10 minutes and further incubated in PBS containing 2.5M Glycine for 5 minutes to stop cross-linking. Cells were collected and resuspended in lysis buffer and sonicated. After centrifugation, the supernatant was diluted in dilution buffer and incubated with Setd2 and H3K36me3 antibodies, respectively, overnight at 4℃. Protein A/G sepharose beads were then added, and the mixture was incubated for an additional 1 hour at 4℃. The sepharose beads were washed twice in low salt wash buffer and then twice in high salt wash buffer, followed by two washes with LiCl buffer and a final two washes with TE buffer. The beads were extracted with 200μl elution buffer and then 8μl NaCl was added into each tube. The mixture was incubated at 65℃ for 4 hours to reverse formaldehyde cross-linking. Additional 1μl (1mg/ml) RNase A was added to each tube and incubated in 37℃ for 30 minutes to remove RNA. Samples were then digested with proteinase K at 50℃ for 4 hours. The purified DNA fragments were subjected to polymerase chain reaction using specific primers. The primers used are listed in Supplementary Table 1.

***RNA-Seq Analysis***

IECs were isolated from Setd2$^{f/f}$ and Setd2$^{Vil\text{-}KO}$ mice treated for 4 d with DSS by EDTA-based isolation. Each sample contained 3 independent repeated animals. NEB Next Ultra Directional RNA Library Prep Kit for Illumina (New England Biolabs, Ipswich, MA, USA) was used for the construction of sequencing libraries. The libraries were then subjected to Illumina sequencing with paired-ends 2x150 as the sequencing mode. The clean reads were mapped to the mouse genome (assembly GRCm38) using the HISAT2 software. Gene expression levels were estimated using FPKM (fragments per kilobase of exon per million fragments mapped) by StringTie. Gene annotation file was retrieved from Ensembl genome browser 90 databases. To annotate genes with gene ontology (GO) terms, Cluster Profiler (R package) was used.

***ChIP-Seq Assay***

IECs were isolated from *Setd2*$^{f/f}$ and *Setd2*$^{Vil\text{-}KO}$ mice after 4 d of DSS treatment. The fragmented chromatin fragments were pre-cleared and then immunoprecipitated with Protein A+G Magnetic beads coupled with anti-H3K36me3 (ab9050) antibody. After reverse crosslinking, ChIP and input DNA fragments were end-repaired and A-tailed using the NEBNext End Repair/dA-Tailing Module (E7442, NEB) followed by adaptor ligation with the NEBNext Ultra Ligation Module (E7445, NEB). The DNA libraries were amplified for 15 cycles and sequenced using Illumina NextSeq 500 with single-end 1x75 as the sequencing mode. Raw reads were filtered to obtain high-quality clean reads by removing sequencing adapters, short reads (length<50 bp) and low-quality reads using Cutadapt (v1.9.1) and Trimmomatic (v0.35). Then FastQC is used to ensure high reads quality. The clean reads were mapped to the mouse genome (assembly GRCm38) using the Bowtie2 (v2.2.6) software. Peak detection was performed using the MACS (v2.1.1) peak finding algorithm with 0.01 set as the p-value cutoff.

***Isolation of Lamina Propria Cells and Flow Cytometry Analysis***

Colons were cut into small pieces and incubated in 5 mM EDTA solution in PBS at 37ºC for 30 min with gentle shaking. After removal of the epithelial layer, the remaining colon pieces were incubated at 37 °C with RPMI medium containing 1.75 mg/ml collagenase A (Roche) and 0.05 mg/ml DNase I (Roche) for 45 min.

After digestion, the supernatant was passed through a 70 μM cell strainer to isolate lamina propria cells. Lamina propria cells were stained for surface markers CD4, CD11b, F4/80 and Gr-1, and subjected to flow cytometry analyses. The antibodies used are listed in Supplementary Table 2.

***Organoid Culture and Analysis***

The intestines were opened longitudinally, and villi were scraped away. The pieces were thorough washed in cold PBS, and incubated in 2 mM EDTA solution in PBS for 10 min at 4 °C. Then, EDTA solution was replaced with PBS and shacked vigorously for 45 s. Crypt fractions were purified by successive centrifugation steps. For every 500-1,000 crypts, a mixture of 100 μL Matrigel (BD Biosciences) and complete growth medium (ratio 2: 1) is added. After polymerization, 100 μl Advanced DMEM/F12 (Invitrogen) containing growth factors (50 ng/ml EGF, PeproTech; 500 ng/ml R-spondin, PeproTech and 100 ng/ml Noggin; PeproTech) was added and refreshed every two or three days. On the fifth day, the organoids were stained with 7-AAD for 5 min, and photos were imaged by Zeiss fluorescence microscope and quantified by Image J software.

***16S-rDNA Sequencing***

Stool DNA samples were analyzed for microbiome at the OE Biotech company. Total genomic DNA was extracted using DNA Extraction Kit following the manufacturer's instructions (or using CTAB methods). Quality and quantity of DNA was verified with NanoDrop and agarose gel. The diluted DNA was used as a template for PCR amplification of bacterial 16S rRNA genes with the barcoded primers and HiFi Hot Start Ready Mix (KAPA). For bacterial diversity analysis, V3-V4 variable regions of 16S rRNA genes was amplified with universal primers 343F and 798R. Clean reads were subjected to primer sequences removal and clustering to generate operational taxonomic units (OTUs) using UPARSE software with 97% similarity cutoff. The representative read of each OTU was selected using QIIME package. All representative reads were annotated and blasted against Silva database Version 123 (or Greengens) (16s rDNA) using RDP classifier (confidence threshold was 70%).

#### Supplemental Figures & Tables

##### Supplemental Figure 1

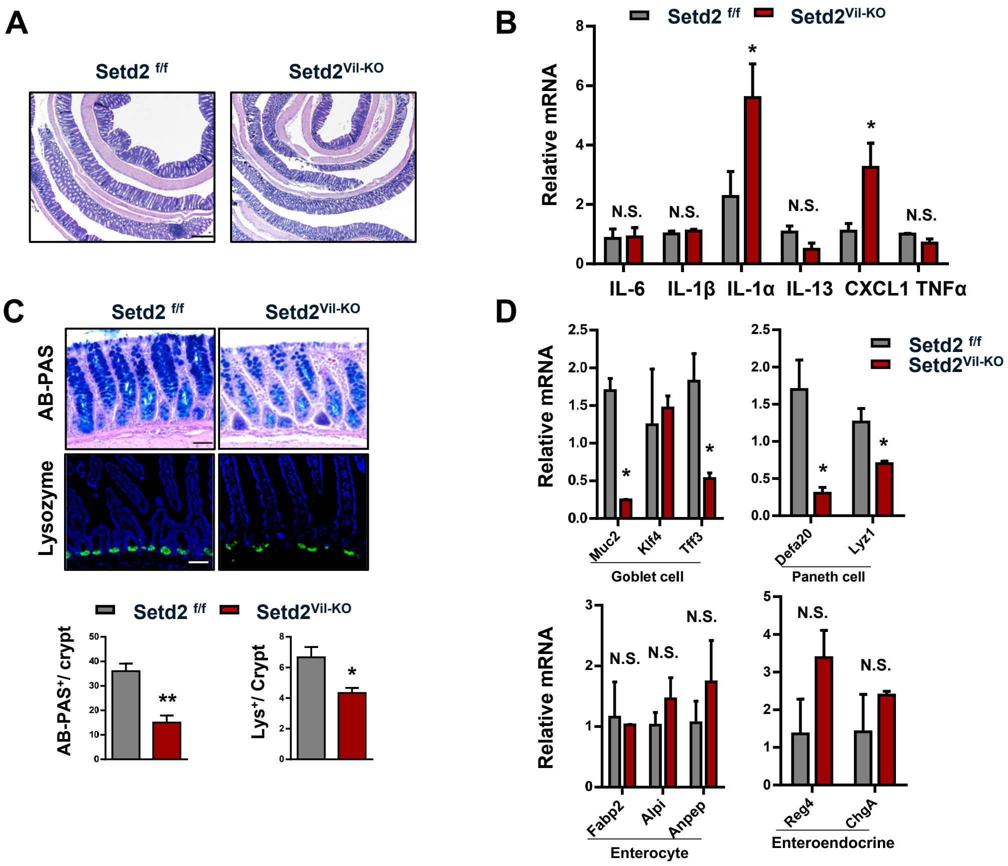


**Supplementary Figure. 1 Deletion of *Setd2* in the colonic epithelium.**

Samples are all derived from 8-week-old *Setd2*$^{Vil\text{-}KO}$ and *Setd2*$^{f/f}$ mice.

**(A)** H&E- stained sections of middle-distal colon tissue as indicated. Scale Bars: 200um.

**(B)** RT-qPCR analysis of whole colon homogenates as indicated.

**(C)** Alcian blue-Periodic acid Schiff (AB-PAS; goblet cells) staining and lysozyme (Lys; Paneth cells) staining in the colon as indicated. Scale Bars: 100um.

**(D)** RT-qPCR analysis of gene expressions in the intestines of *Setd2*$^{Vil\text{-}KO}$ and *Setd2*$^{f/f}$ mice.

The data represent the mean ± S.E.M, and statistical significance was determined by a two-tailed Student's t-test. * $p < 0.05$, ** $p < 0.01$. N.S., Not Significant.

#### Supplemental Figure 2

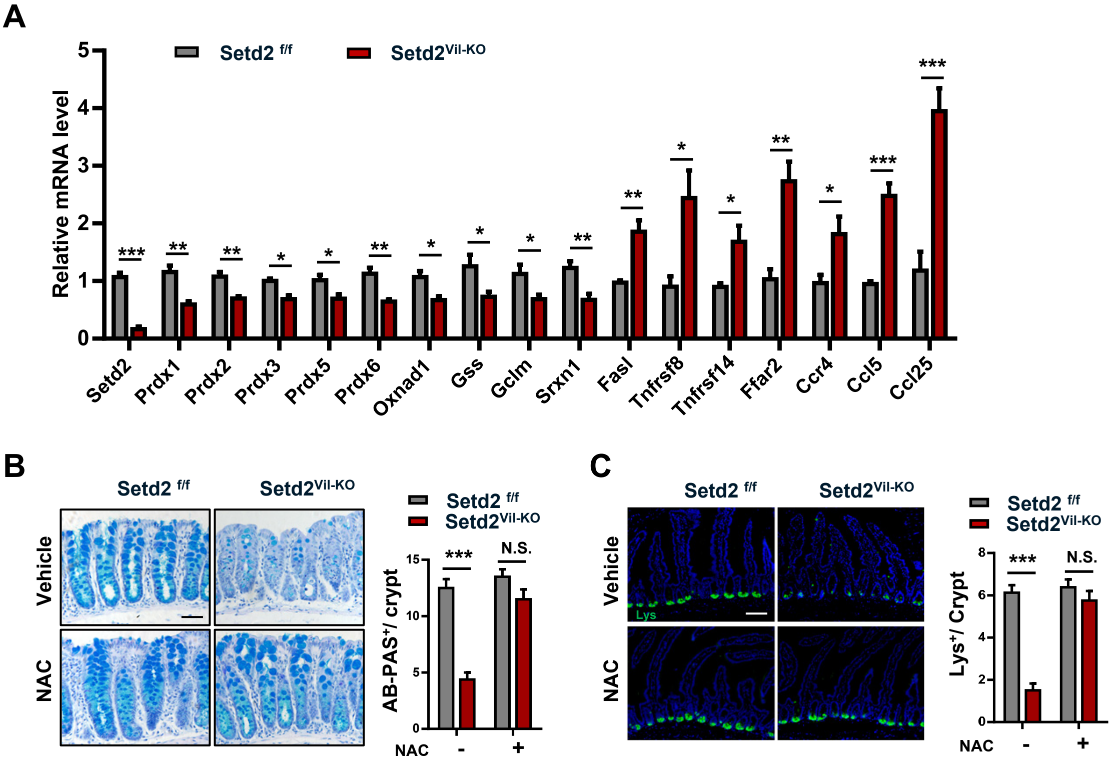


**Supplementary Figure. 2 Increased ROS leads to colitis in *Setd2*$^{Vil\text{-}KO}$ mice.**

**(A)** Relative expression of differential genes in IECs from DSS-treated *Setd2*$^{Vil\text{-}KO}$ and *Setd2*$^{f/f}$ mice.

**(B, C)** Alcian blue-Periodic acid Schiff (AB-PAS; goblet cells) staining **(B)** and lysozyme (Lys; Paneth cells) staining **(C)** of DSS-treated *Setd2*$^{Vil\text{-}KO}$ and *Setd2*$^{f/f}$ mice with or without NAC treatment. Scale Bars: 100um.

The data represent the mean ± S.E.M, and statistical significance was determined by a two-tailed Student‘s t-test. * $p < 0.05$, ** $p < 0.01$, *** $p < 0.001$. N.S., Not Significant.

### Supplemental Figure 3

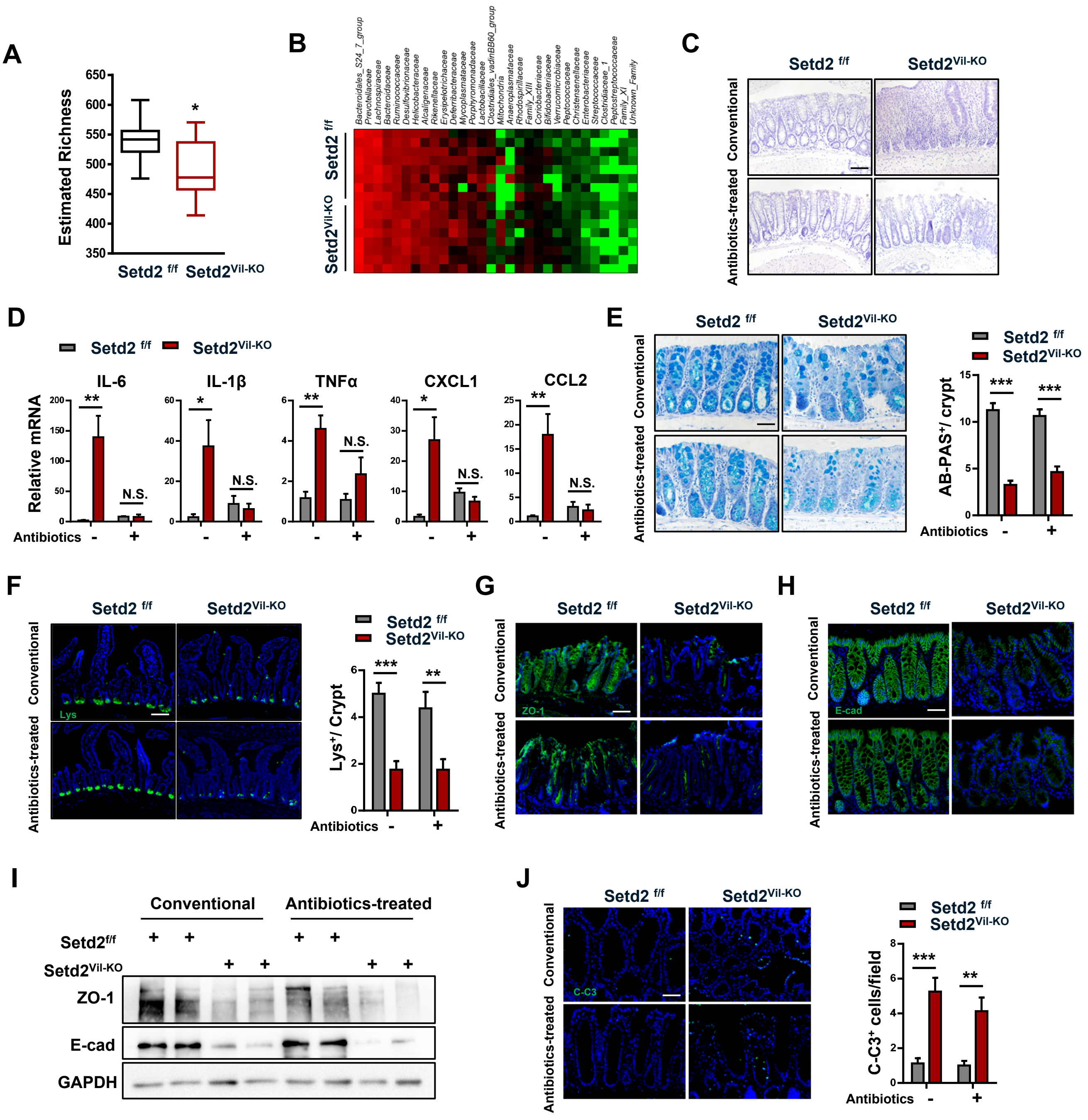

**Supplementary Figure. 3 Analyses in the *Setd2*$^{Vil-KO}$ and *Setd2*$^{f/f}$ mice with or without antibiotics treatment.**

(A) Chao bacterial diversity in *Setd2*$^{Vil-KO}$ and *Setd2*$^{f/f}$ mice.

(B) Heat map of bacterial family in the intestinal microbiota of *Setd2*$^{Vil-KO}$ and *Setd2*$^{f/f}$ mice.

(C) H&E staining in colon sections as indicated. Scale Bars: 100um.

**(D)** RT-qPCR analysis of colon homogenates from *Setd2*$^{Vil-KO}$ and *Setd2*$^{f/f}$ mice to assess cytokine and chemokine productions.

**(E, F)** Alcian blue-Periodic acid Schiff (AB-PAS; goblet cells) staining **(E)** and lysozyme (Lys; Paneth cells) staining **(F)** in colon sections as indicated, and quantitation results are shown in the right (n=8). Scale Bars: 100um.

**(G, H)** ZO-1 staining **(G)** and E-cadherin staining **(H)** in colon sections as indicated. Scale Bars: 100um.

**(I)** Colon lysates were analyzed by western blotting with the indicated antibodies.

**(J)** Cleaved caspase-3 staining as indicated, and quantitation results are shown in the right (n=8). Scale Bars: 50um.

The data represent the mean ± S.E.M, and statistical significance was determined by a two-tailed Student‘s t-test. * $p < 0.05$, ** $p < 0.01$, *** $p < 0.001$. N.S., Not Significant.

**Supplementary Table 1: Primers for RT-qPCR, ChIP-qPCR analysis and genotyping**

**Primers for RT-qPCR**

| Gene name | Sense 5’-3’ | Antisense 5’-3’ |
|---|---|---|
| hGapdh | AGAAGGCTGGGGCTCATTTG | AGGGGCCATCCACAGTCTTC |
| hSetd2 | GAACCCTTACCGGAAACCTGA | CAGGTCCTCAGGATTCTTACAG |
| mGapdh | AGGTCGGTGTGAACGGATTTG | TGTAGACCATGTAGTTGAGGTCA |
| mSetd2 | CATAGCTGTGAACCAAACTGTGA | TAATTCTGAGCCTGAAGGAACTA |
| mIL-6 | AGTTGCCTTCTTGGGACTGA | CAGAATTGCCATTGCACAAC |
| mIL-13 | AAGGAGCTTATTGAGGAGCTG | TCAGGGAATCCAGGGCTACA |
| mIL-1α | GAGAGCCGGGTGACAGTATC | TGACAAACTTCTGCCTGACG |
| mIL-1β | GAAATGCCACCTTTTGACAGTG | TGGATGCTCTCATCAGGACAG |
| mIL-10 | TAGAGCTGCGGACTGCCTTC | CATTTCCGATAAGGCTTGGCAA |
| mTNF-α | CGTCAGCCGATTTGCTATCT | CGGACTCCGCAAAGTCTAAG |
| mCXCL1 | CTGGGATTCACCTCAAGAACATC | CAGGGTCAAGGCAAGCCTC |
| mCCL2 | TTAAAAACCTGGATCGGAACCAA | GCATTAGCTTCAGATTTACGGGT |
| mMuc2 | GCCCGTGGAGTCGTACGTGC | TTGGGGCAGAGTGAGGCGGT |
| mKlf4 | GGCGAGTCTGACATGGCTG | GCTGGACGCAGTGTCTTCTC |
| mTff3 | TAATGCTGTTGGTGGTCCTG | CAGCCACGGTTGTTACACTG |
| mRega4 | GGCGTGCGGCTACTCTTAC | GAAGTACCCATAGCAGTGGGA |
| mChgA | AAGTGCGTCCTGGAAGTCATCTC | GCTTGGCTTTTCTGGCTTGC |
| mDefa20 | TGTAGAAAAGGAGGCTGCAATAG | AGAACAAAAGTCGTCCTGAGC |
| mLyz1 | GAGACCGAAGCACCGACTATG | CGGTTTTGACATTGTGTTCGC |
| mFabp2 | GTGGAAAGTAGACCGGAACGA | CCATCCTGTGTGATTGTCAGTT |
| mAlpi | GGCCATCTAGGACCGGAGA | TGTCCACGTTGTATGTCTTGG |
| mAnpep | ACGCTCAGGAGAAGAATAGGAA | CTTAGGCAAGCGATACTGGTTC |
| mPrdx1 | AATGCAAAAATTGGGTATCCTGC | CCGTGGGACACACAAAAGTAAA |
| mPrdx2 | GATGGTGCCTTCAAGGAAATCA | CCGTGGGGCAAACAAAAGTG |
| mPrdx3 | GGTTGCTCGTCATGCAAGTG | CCACAGTATGTCTGTCAAACAGG |
| mPrdx5 | GCTGCAAAGCCAGTTCTGTG | CCACTGAGGGAATGGCATCTC |
| mPrdx6 | CGCCAGAGTTTGCCAAGAG | TCCGTGGGTGTTTCACCATTG |
| mOxnad1 | GCCCACGTTGCCTTGATTC | AGGGTAAGGTGACGCAAAGTG |
| mGss | AAAGCAGGCCATAGACAGGG | TGAATGGGGCATACGTCACC |
| mGclm | AGGAGCTTCGGGACTGTATCC | GGAAACTCCCTGACTAAATCGG |
| mSrxn1 | CCCAGGGTGGCGACTACTA | GTGGACCTCACGAGCTTGG |
| mFasl | TCCGTGAGTTCACCAACCAAA | GGGGGTTCCCTGTTAAATGGG |
| mTnfrsf8 | CCTTCCCAACGGATCGACC | CCCGTCTTCATTGACGTAGTAGT |
| mTnfrsf14 | CAGGCCCCTACAGACAACAC | ACTCGTCTCCCACAAGGAACT |
| mFfar2 | CTTGATCCTCACGGCCTACAT | CCAGGGTCAGATTAAGCAGGAG |
| mCCR4 | TGCACCAAGGAAGGTATCAAGG | GTACACGTCCGTCATGGACTT |
| mCCL5 | GCTGCTTTGCCTACCTCTCC | TCGAGTGACAAACACGACTGC |
| mCCL25 | GACTGCTGCCTGGGTTACC | TGGCGGAAGTAGAATCTCACA |

**Primers for ChIP-qPCR**

| Gene name | Sense 5'-3' | Antisense 5'-3' |
|---|---|---|
| Prdx3 | GATATGGCTCACCTGGTA | TTAGCTCAGATGACTTCTG |
| Prdx6 | GGCTATAGCCCAATGCCCAC | AAAGTATTTACAAGTATTTGGA |
| Gclm | TCGTCCCCTGACTCTTGCTC | TGAAAGGTCTGAGGAAGCAGCT |
| Srxn1 | TGGAGTCCTTCCCGCCTCAAG | GGGTTTTAAGGTTCCCCAAATT |

**Primers for genotyping**

| Gene name | Sense 5'-3' | Antisense 5'-3' |
|---|---|---|
| Villin-Cre | GTGTTTGGTTTGGTTTCCTCTGCATAAGA | GCAGGCAAATTTTGGTGTACGGTCA |
| Setd2 | GTAAAGTAGTATTATGCCAAGGCCC | TATTTAAACTCTCTCTGGGGGTGG |

**Supplementary Table 2: Antibodies**

**Antibodies used for flow cytometry**

| ID | Clone/Catalog NO. | Vol./$10^6$ cells | Brand |
|---|---|---|---|
| CD4 | G/C1.5 | 1.5uL | eBioscience™ |
| CD11b | M1/70 | 1.5uL | eBioscience™ |
| F4/80 | BM8 | 1.5uL | eBioscience™ |
| Gr-1 | RB6-8C5 | 1.5uL | eBioscience™ |
| H2DCFDA | C6827 | 5mM | Life Technologies |

**Antibodies used for IHC(IF) and western blotting**

| ID | Clone/Catalog NO. | Dilution | Brand |
|---|---|---|---|
| SETD2 | C332416 | 1:1000 | LSBio |
| H3K36me3 | ab9050 | 1:1000 | Abcam |
| H3 | ab10799 | 1:1000 | Abcam |
| Lysozyme | A0099 | 1:500 | Dako |
| F4/80 | BM8 | 1:500 | eBioscience™ |
| Ki67 | ab15580 | 1:2000 | Abcam |
| ZO-1 | 61-7300 | 1:1000 | Thermo |
| E-cad | #3195T | 1:500 | Cell Signaling Technology |
| Claudin-1 | # 4933T | 1:1000 | Cell Signaling Technology |
| Caspase3 | #9662 | 1:1000 | Cell Signaling Technology |
| C-Caspase3 | #9661 | 1:1000 | Cell Signaling Technology |
| 8-OHdG | # ab48508 | 1:500 | Abcam |
| PRDX6 | ab59543 | 1:1000 | Abcam |
| GAPDH | #5174 | 1:1000 | Cell Signaling Technology |
